# An antibiotic-free antimicrobial combination of bacteriocins and a peptidoglycan hydrolase: *in vitro* and *in vivo* assessment of its efficacy

**DOI:** 10.1101/2024.11.19.624290

**Authors:** Christian Kranjec, Thomas F. Oftedal, Kirill V. Ovchinnikov, Vinicius da Silva Duarte, Simen Hermansen, Magdalena Kaus-Drobek, Izabela Sabała, Davide Porcellato, Harald Carlsen, Morten Kjos

**Affiliations:** Faculty of Chemistry, Biotechnology and Food Science, Norwegian University of Life Sciences, Ås, Norway; Department of Molecular Oncology, Institute for Cancer Research, Oslo University Hospital, Oslo, Norway; Mossakowski Medical Research Institute Polish Academy of Sciences, Warsaw, Poland

## Abstract

Mastitis is an inflammatory disease of the mammary gland commonly brought about by bac-terial pathogens that gain physical access to the glandular epithelium through the teat canal. In bovines, common mastitis-causing agents are environmental or pathogenic bacterial spe-cies, including staphylococci, streptococci, enterococci, and Gram-negative bacteria such as *Escherichia coli*. Current therapeutic strategies for bovine mastitis typically involve the ad-ministration of antibiotic formulations within the infected udder, possibly resulting in in-creased selection of antibiotic resistance and the accumulation of antibiotic residues within the milk. In this study, we sought to design an antibiotic-free antimicrobial formulation to treat bovine mastitis based on bacterial antimicrobial peptides (bacteriocins) and proteins (pepti-doglycan hydrolases). Using a combination of *in vitro* assays with a range of bacteriocins, we show that the combination of the thiopeptide micrococcin P1 (MP1) and the lantibiotic nisin A (NisA) is a robust antimicrobial formulation that effectively inhibits the growth of bo-vine mastitis-derived bacteria, both in planktonic and biofilm-associated growth modes. The addition of AuresinePlus (Aur, a staphylococcus-specific PGH) further increased the antimi-crobial potency against *S. aureus*. Furthermore, using two mouse models, a skin infection model and a mastitis model, we show that the combination MP1-NisA-Aur effectively inhibits methicillin-resistant *S. aureus* (MRSA) *in vivo*. We discuss the potential and challenges of using antibiotic-free antimicrobial combinations in the treatment of bacterial infections.

## Introduction

Mastitis is a condition caused by the inflammation of the mammary gland, most commonly brought about by bacterial infections but also by physical trauma of the udder. In the dairy cattle industry, mastitis is a serious concern for the welfare of the animals and an economic burden for farmers. It has been estimated that mastitis result in a gross margin loss of 11 to 18% per cow per year, with 70% of the economic burden related to mammary tissue damage leading to reduced milk production.^1,2^ Bacterial intra-mammary infections (IMIs), the main cause of bovine mastitis, can result from contagious mastitis-causing bacteria (i.e., *Staphylococcus aureus*, *Streptococcus agalactiae, Mycoplasma* spp.) or from environmental bacteria (i.e., coliform bacteria, *Enterococcus* spp., coagulase-negative, non-*aureus* staphylococci, *Streptococcus uberis*).^3–7^ Mastitis can be classified as clinical or sub-clinical, where the former is characterized by overt signs of inflammatory disease and a reduction in milk production and quality. Subclinical mastitis is manifested by reduced milk production and increased somatic cell count (SCC) in the milk without overt signs of disease, which is more common in older lactating animals.^8–10^ Subclinical mastitis also poses increased public health concerns linked to the consumption of dairy products contaminated by pathogenic agents or their thermostable toxins.^11,12^

Current mastitis treatments involve the use of antibiotics administered via intra-mammary infusion, intramuscular, or intravenous injections.^13^ Antibiotic therapy, however, poses serious concerns due to the high rate of therapeutic failure and accumulation of the antibiotic in the milk. Furthermore, due to the widespread use of antibiotics, infections are more frequently caused by antibiotic-resistant strains, which renders antibiotic treatments ineffective.^14^ In addition, some bacterial species, such as *Staphylococcus aureus*, have a marked capacity to form biofilms, which strongly enhances their resilience to antibiotic therapy.^15^ The timing of initiating mastitis therapy is crucial; the best therapeutic results are obtained upon administration of the antimicrobial therapy during the dry period; the 6-8 weeks prior to calving where the cow is not milked.^16,17^ Additionally, treatment of IMI during the dry period (dry cow therapy) prevents the production of nonsalable waste milk.^18^ Potential alternative antimicrobial therapies include the use of prophylactic vaccines, bacteriophages, bacterially-derived antimicrobial agents such as bacteriocins and bacteriolytic enzymes.^19–22^

Bacteriocins, ribosomally-synthesized bacterial peptides, are prime examples of metabolites with antimicrobial activity. It is thought that bacteriocins are produced by virtually all bacteria and, from an ecological point of view, are synthesized in order to confer a selective advantage to the producer in niche competition, as these molecules often display activity against closely related species.^23–25^ Bacteriocins generally kill target cells by disruption of the cell membrane (e.g., nisin, garvicin KS, enterocin EJ97),^26,27^ but there are also bacteriocins with intracellular targets, such as micrococcin P1 which inhibits translation.^28^ Bacteriocins that are post-translationally modified, such as nisin and micrococcin P1, are typically regarded as belonging to class I; while bacteriocins that are unmodified, which includes garvicin KS and enterocin EJ97 belong to class II.^23^ In the last decades, bacteriocins produced by lactic acid bacteria (LAB) have received greater attention for their therapeutic potential.^22^ LAB are an essential part of many dairy products, therefore LAB and the metabolites they produce are generally recognized as safe (GRAS) according to the Food and Drug Administration (FDA). With respect to this, the well characterized LAB bacteriocins nisin and pediocin PA-1 have been approved for use as food preservatives.^25^ It is also generally accepted that LAB bacteriocins have potential medical applications since they inhibit growth of important human pathogens, including *S. aureus* MRSA, vancomycin-resistant enterococci (VRE) and *Listeria monocytogenes*, and are effective against biofilm-forming strains.^23,29–34^ Furthermore, since LAB are naturally present in raw dairy milk they can potentially be used as probiotics to prevent and treat bovine mastitis.^35–39^

Peptidoglycan hydrolases (PGHs), enzymes that catalytically degrade the peptidoglycan (PG) layer of the bacterial cell wall, have also been explored as potential alternatives to antibiotics.^40–44^ These hydrolytic enzymes can originate from bacteria (autolysins and exolysins) or bacteriophages (endolysins).^20,43,45^ Peptidoglycan hydrolases are often further modified to increase their potency, stability or specificity by domain shuffling and/or site-directed mutagenesis. These modifications result in enzymes that are more effective under a broader range of conditions which extends their potential applications.^40,41,46^ Notably, similar to bacteriocins from LAB, PGHs exhibit high target specificity for prokaryotic cells, which is thought to make them a safe option for antimicrobial treatment in human and veterinary medicine. Moreover, they display a low prevalence of resistance, a feature particularly relevant in the era of antimicrobial resistance.^47^

In this study, we explored the potential use of an antibiotic-free and bacteriocin-based combination against a broad collection of Gram-positive mastitis-derived bacterial strains. We show that the combination between nisin A and the thiopeptide bacteriocin micrococcin P1 serves as a scaffold to further build antibiotic-free antimicrobial combinations with increased potency and reduced occurrence of resistance. When combined with a PGH with high specificity to *Staphylococcus* spp., AuresinePlus, the combination showed to have a superior activity against biofilms formed by a panel of mastitis-derived staphylococci, including *S. aureus*. In addition, we show that the tricomponent combination is effectively eradicated infections caused by methicillin-resistant *S. aureus* (MRSA) in two murine models.

## Results

### A combination of micrococcin P1, nisin A and AuresinePlus is highly effective against a wide array of mastitis-derived strains

Natural antimicrobial proteins and peptides may become a powerful tool against bacterial infections to tackle the problem of antibiotic resistance. Here, we investigated the possibility of using bacteriocins and AuresinePlus (Aur), a *Staphylococcus*-specific chimeric peptidoglycan hydrolase (PGH), to design an antibiotic-free combination to treat bovine mastitis.^48^ We assessed the activity of four bacteriocins with characterized spectra of activity: micrococcin P1 (MP1), garvicin KS (GarKS), EntEJ97-short (EJs) and nisin A (Nis) against a pool of representative mastitis-derived bacterial species belonging to the genera *Staphylococcus*, *Streptococcus*, *Enterococcus* and *Trueperella.*^49,50,37,51–53^ Antimicrobials such as bacteriocins can significantly reduce bacterial populations, but often fail to provide long-term inhibition due to resistance development, proteolytic degradation, persister cells or other detoxification mechanisms in bacteria. As such we assessed the ability of the combinations to provide long-term inhibition for up to 1 week (168 h). As shown in Figures S1A and S1B, these assays identified bacteriocin combinations containing MP1 and Nis (MP1/Nis, MP1/Nis/EJs or MP1/Nis/GarKS) as able to provide a durable antimicrobial activity against planktonic mastitis-derived isolates in terms of low minimal inhibitory concentration (MIC). Next, we were interested in verifying the effectiveness of the MP1/Nis-containing antimicrobial combinations against a wider panel of 116 bovine mastitis-derived Gram-positive isolates (Figure S2A). The effectiveness of the antimicrobial combination was determined by measuring the minimum inhibitory concentration (MIC) of planktonic cells at 5, 24, 48 and 168 h after addition of the antimicrobials to cultures, the results are presented in Figure 1 and Figure S2B. As can be seen in Figure 1, while the MP1/Nis combination effectively inhibited planktonic growth of all strains, the addition of either EJs or GarKS resulted in a further reduction in MIC for all the bacterial strains (indicated by a shift in the blue vertical MIC line towards the left from panel A to B and C).

**Figure 1.**
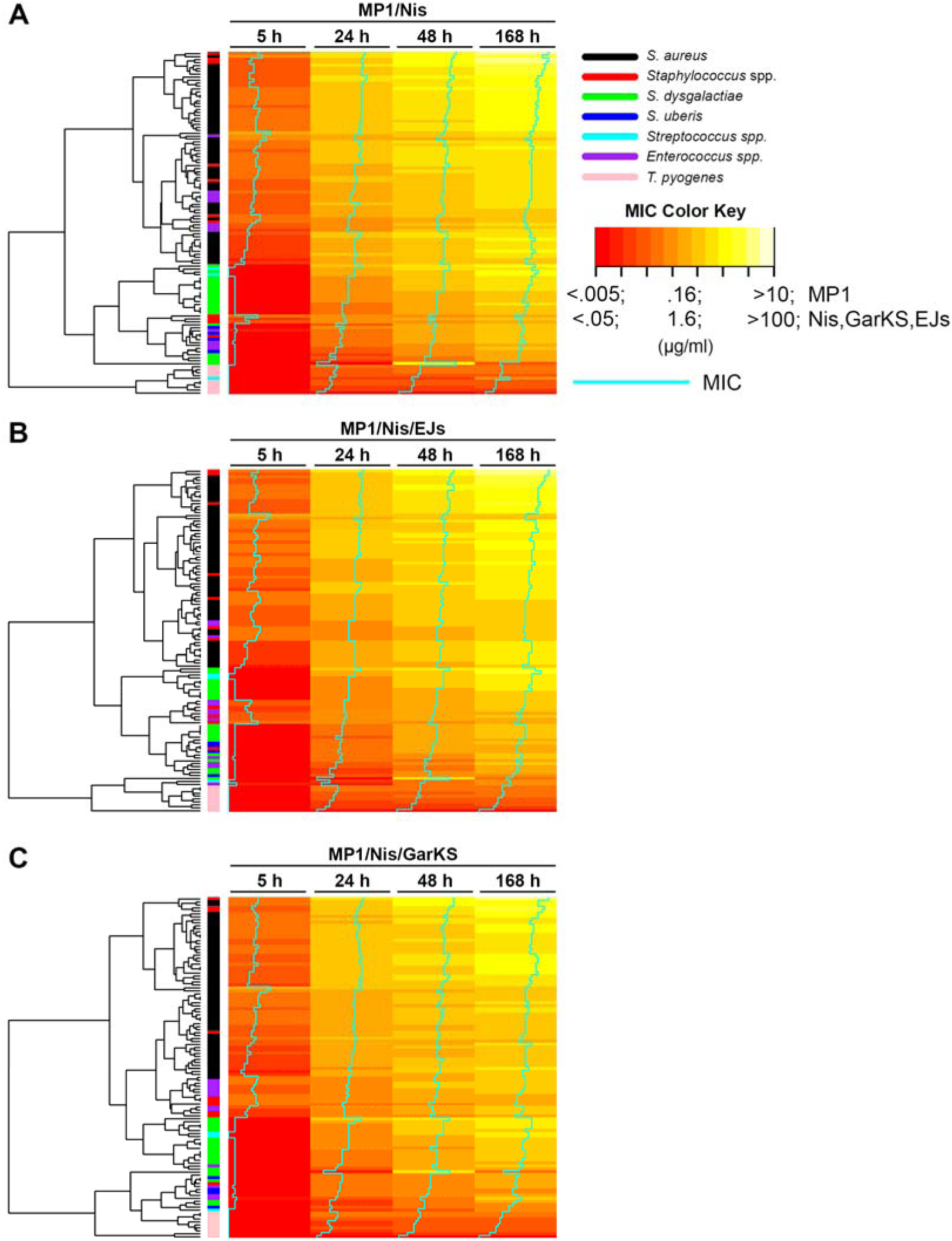
Heatmap showing the distribution of MIC values (µg/ml) for the indicated mastitis-derived species against the antimicrobial combinations (a) MP1/Nis, (b) MP1/Nis/EJs, and (c) MP1/Nis/GarKS. MIC values were measured 5, 24, 48 and 168 h after addition of the antimicrobial combinations. MIC values are indicated by color from red (low) to yellow-white (high) as indicated on the MIC color key and by a vertical line (in cyan) going from low (left) to high (right). The species of the strains tested (n = 116) are indicated by color (see legend). A dendrogram showing the clustering of strains based on their MIC-values is shown to the left.

Staphylococci (*S. aureus* and *Staphylococcus* spp.) were however more resilient resilient with some strains displaying consistently high MIC for all the treatment regimes. *Trueperella pyogenes* displayed the lowest MICs among the strains in the panel (Figure 1 and Figure S2B). The addition of AuresinePlus to the MP1/Nis combination led to a highly significant decrease of the MIC for all *S. aureus* isolates, irrespective of the treatment duration, and reduced the inter-strain variability within the *S. aureus* cohort (Figure 2 and Figure S3).

**Figure 2.**
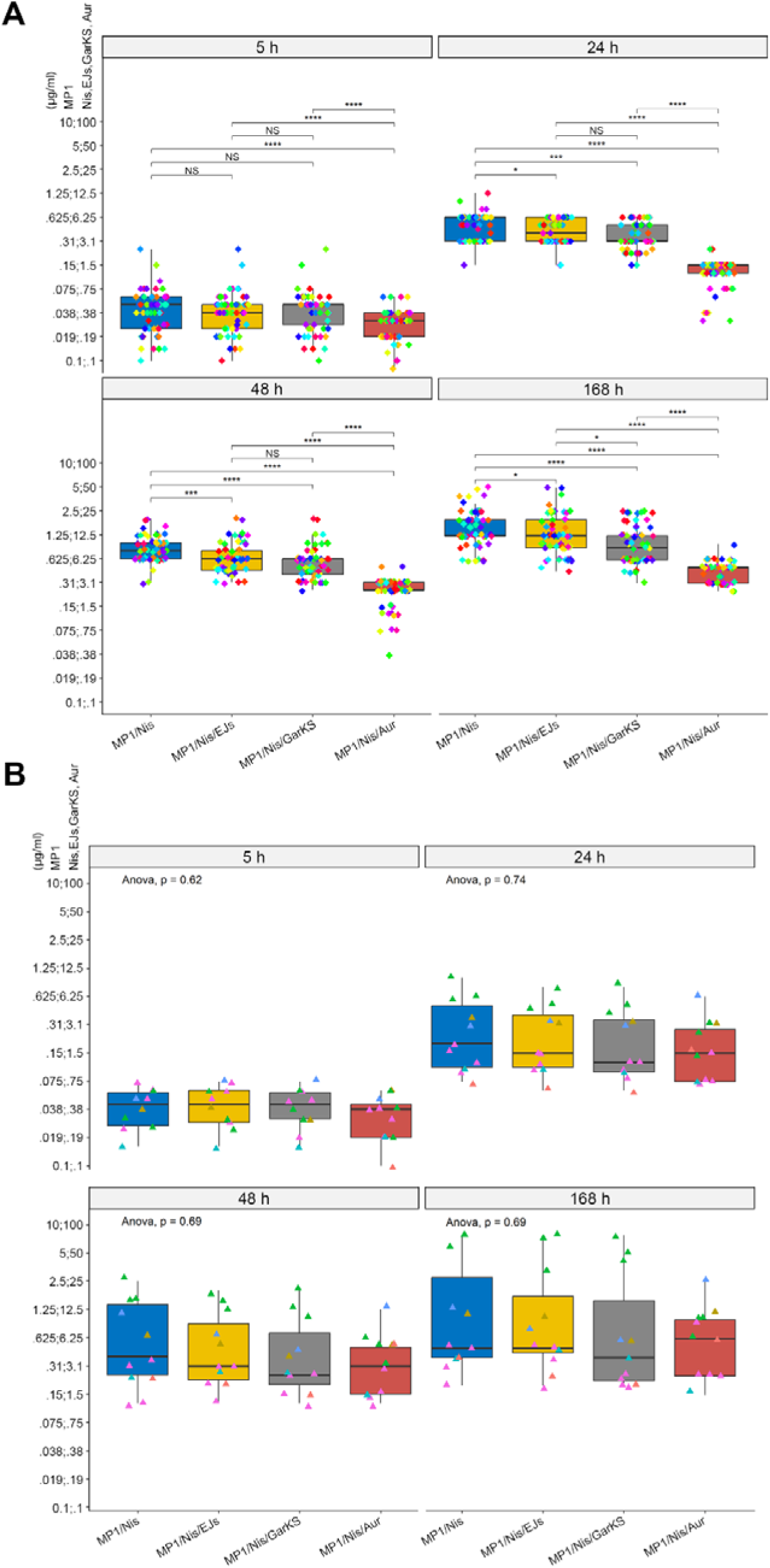
Aur potentiates antimicrobial effect of bacteriocins against mastitis-derived *S. aureus*. (a) Box and whisker plot (Tukey’s method) showing the distribution in MIC for 57 strains of *S. aureus* included in our mastitis collection against the MP1/Nis, MP1/Nis/EJs, MP1/Nis/GarKS or MP1/Nis/Aur combinations, measured 5, 24, 48 or 168 h after addition of the antimicrobial combination. (b) Same as in panel A, but against 11 strains of non-*aureus* staphylococcal species. In panel A, post-hoc statistical analysis was performed as pairwise comparisons with the Wilcoxon test, asterisk representation of statistical significance: *p ≤ 0.05; ***p ≤ 0.001; **** p ≤ 0.0001; ns = not significant. In panel B, the global statistical significance was assessed using the one-way ANOVA test (p-values indicated).

Conversely, in line with the known specificity of AuresinePlus, the contribution of the hydrolase against other staphylococcal species was less obvious; with only some of the isolates within the *Staphylococcus* spp. cohort (e.g. *S. simulans* and *S. haemolyticus*) being more susceptible to the hydrolase-containing combination (Figure 2 and Figure S3).^41^ Taken together, these results indicate that the antimicrobial spectrum of activity of the MP1/Nis/Aur combination covers the most important Gram-positive mastitis-associated pathogens.

### Bacteriocins and AuresinePlus retain activity against biofilms of mastitis-derived bacteria

We next assessed whether our antimicrobial combinations could eradicate cells in biofilms produced by mastitis-associated bacteria, since biofilm formation has been associated with persistence of mastitis infections and potentially be responsible for cases of chronic mastitis.^54,55^ By performing biofilm formation assays *in vitro*, we found that 92% of *Staphylococcus* strains and 58% of *Enterococcus* strains were good or strong biofilm formers (Figure S4A-S4C), whereas the streptococcal isolates were mostly weak biofilm formers (Figure S4D). Only weak or no biofilm formation was found among *T. pyogenes* isolates (data not shown).

A modified version of the biofilm-oriented antimicrobial test (BOAT) was used to test whether biofilms formed by *S. aureus*, *Staphylococcus* spp. and enterococci were sensitive to combinations of bacteriocins and PGH. In this assay, the metabolic activity indicator triphenyl tetrazolium chloride is used to indicate metabolic activity (viability) of biofilm-associated cells after antimicrobial treatment.^32,34,51,56,57^ As can be seen in Figure 3, both MP1/Nis and MP1/Nis/Aur combinations significantly reduced biofilm-associated metabolic activity in 12 mastitis-associated strains of *S. aureus* after 24 h (Figure 3A and 3B).

**Figure 3.**
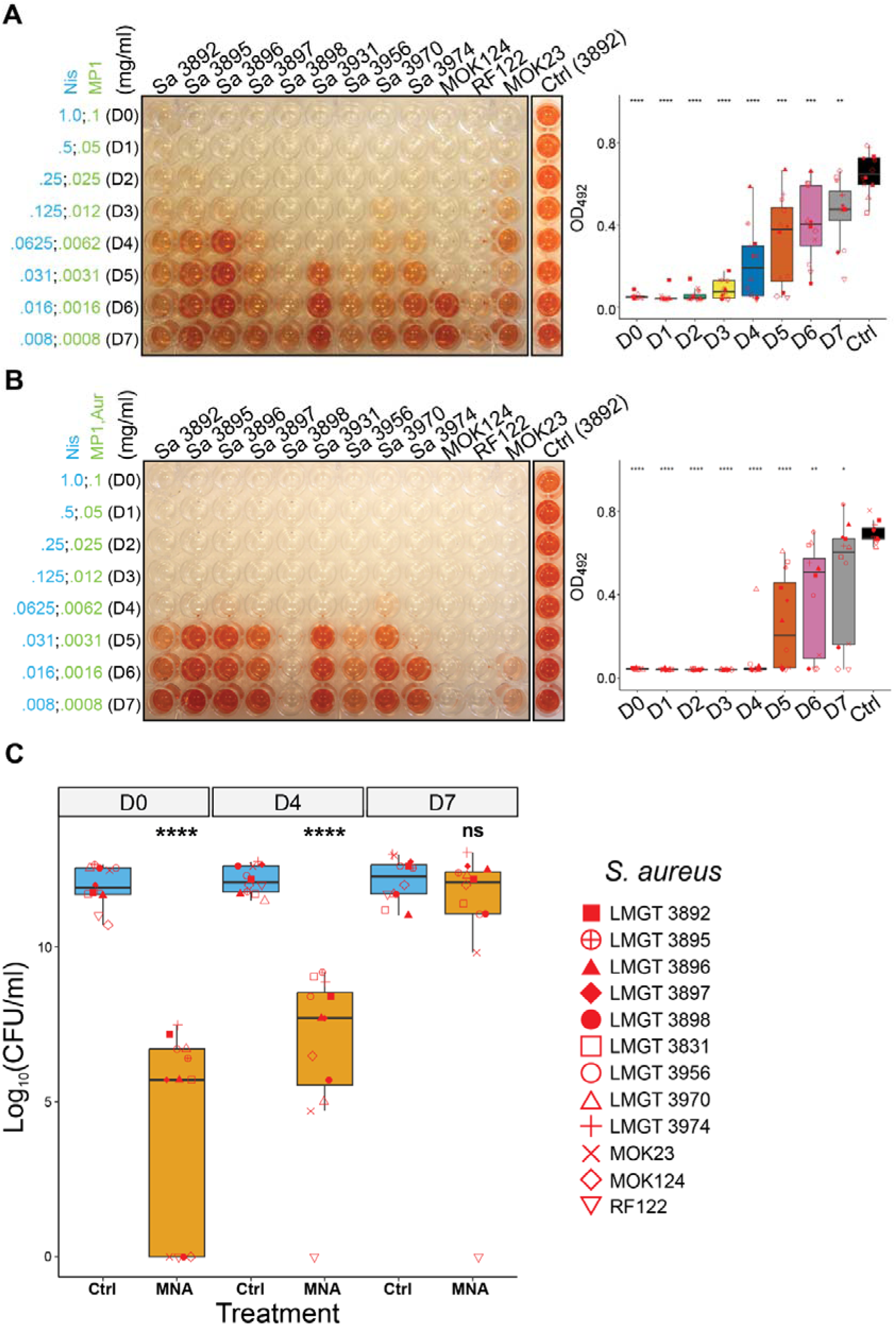
Assessment of bacteriocin and bacteriocin/AuresinePlus combinations against a panel of mastitis-derived *S. aureus* strains. (a) The left panel shows a representative image of the BOAT assay performed with a two-fold dilution of the MP1/Nis combination against the indicated *S. aureus* strains. The concentrations of the antimicrobials (in mg/ml) are shown on the left (dilutions D0-D7). The assays were simultaneously performed with the control vehicles of the antimicrobials (Ctrl), and a representative image of *S. aureus* strain 3892 is shown. The amount of the red color is an indirect measure of metabolic activity and was measured by optical density readings at 492 nm (OD_492_). A plot of the quantification of red color (bacterial metabolic activity) as a function of the dilution factor is shown in the right panel. (b) Same as in panel A except the strains were tested against the MP1/Nis/Aur combination. Statistical analyses of each group (D0-D7) relative to the vehicle control (Ctrl) was performed with Welch’s t-test. (c) Boxplot showing the median distribution of log10-transformed colony forming units (Log_10_CFU/ml) calculated after the BOAT assay for the same strains as in panels A and B. The CFU quantification was performed for the MP1/Nis/Aur combination (MNA) at concentrations D0, D4 and D7 depicted in panel B. Asterisk representation of statistical significance: *p ≤ 0.05; **p ≤ 0.01; ***p ≤ 0.001; **** p ≤ 0.0001. Statistical plots (box and whisker) according to Tukey’s method.

Notably, the susceptibility of all strains to the antimicrobials was reduced (ranging from 2.5-to 80-fold) compared to planktonic cells. The addition of AuresinePlus to the combination significantly reduced metabolic activities compared to MP1/Nis alone (*p* < 0.05, Figure 3A, 3B). When assessing cell viability by determining colony forming units (CFU) (Figure 3C), a very significant overall reduction could be observed for MP1/Nis/Aur-treated *S. aureus* biofilms (*p* < 0.0001; Figure 3C). Although metabolic activity measurements indicated no-or low cell viability in the biofilms after treatment at D0-D4, seven out of twelve strains were shown to viable after treatment at D0 by CFU counting (CFU/ml > 10^5^), and eleven out of twelve at D4. Such discrepancy between CFU data and metabolic activity has also been observed previously.^31^ The highest sensitivity to the MP1/Nis/Aur combination was seen for strain RF112, for which neither metabolic activity nor cell viability could be detected even at the lowest concentration tested (Figure 3). Among other mastitis-derived species, biofilm-associated enterococci were highly sensitive to the MP1/Nis combinations (Figures S5A). Against non-*aureus* staphylococcal species, AuresinePlus extended the overall effective concentration window of the antimicrobial combination (Figure S6A and S6B) and, consequently, reduced the overall cell viability (Figure S6C).

Taken together these data indicate that the bacteriocins and AuresinePlus make an effective antimicrobial combination active on biofilm-associated bacterial cells to reduce their viability.

### Effect of the combination of micrococcin P1, nisin A and AuresinePlus against *S. aureus* in a murine skin infection model

To assess the therapeutic potential of the MP1/Nis/PGH combination *in vivo*, we utilized a murine skin infection model together with a bioluminescent methicillin-resistant *S. aureus* (MRSA) strain Xen31 (Perkin Elmer, bioluminescent derivative of ATCC 33591). The skin infection model has previously been used and validated in our laboratory for the *in vivo* study of different antimicrobial combinations.^30,31^ Furthermore, *S. aureus* Xen31 exhibit comparable susceptibility to the MP1/Nis/Aur combination in *in vitro* assays as the mastitis-associated strains used in this study (Figure S7A and S7B).

Skin infections were established on the back skin of female BALB/c mice by inoculating 10^7^ CFU of *S. aureus* Xen31 in freshly produced wounds of 10 mm in diameter. The infection was allowed to establish for 24 h prior to initiating antimicrobial treatment at day 1 post infection (PI). Importantly, at this time point infections had established and the bacterially-produced bioluminescence could be detected (Figure S8). Mice were randomly divided into two groups: group 1 (MP1/Nis/Aur-treated) and group 2 (vehicle-treated controls). Mice were then subjected to bioluminscence imaging daily from day 1 PI to day 7 PI, while treatment was administered daily from 1 through 5 PI (Figure 4A).

**Figure 4.**
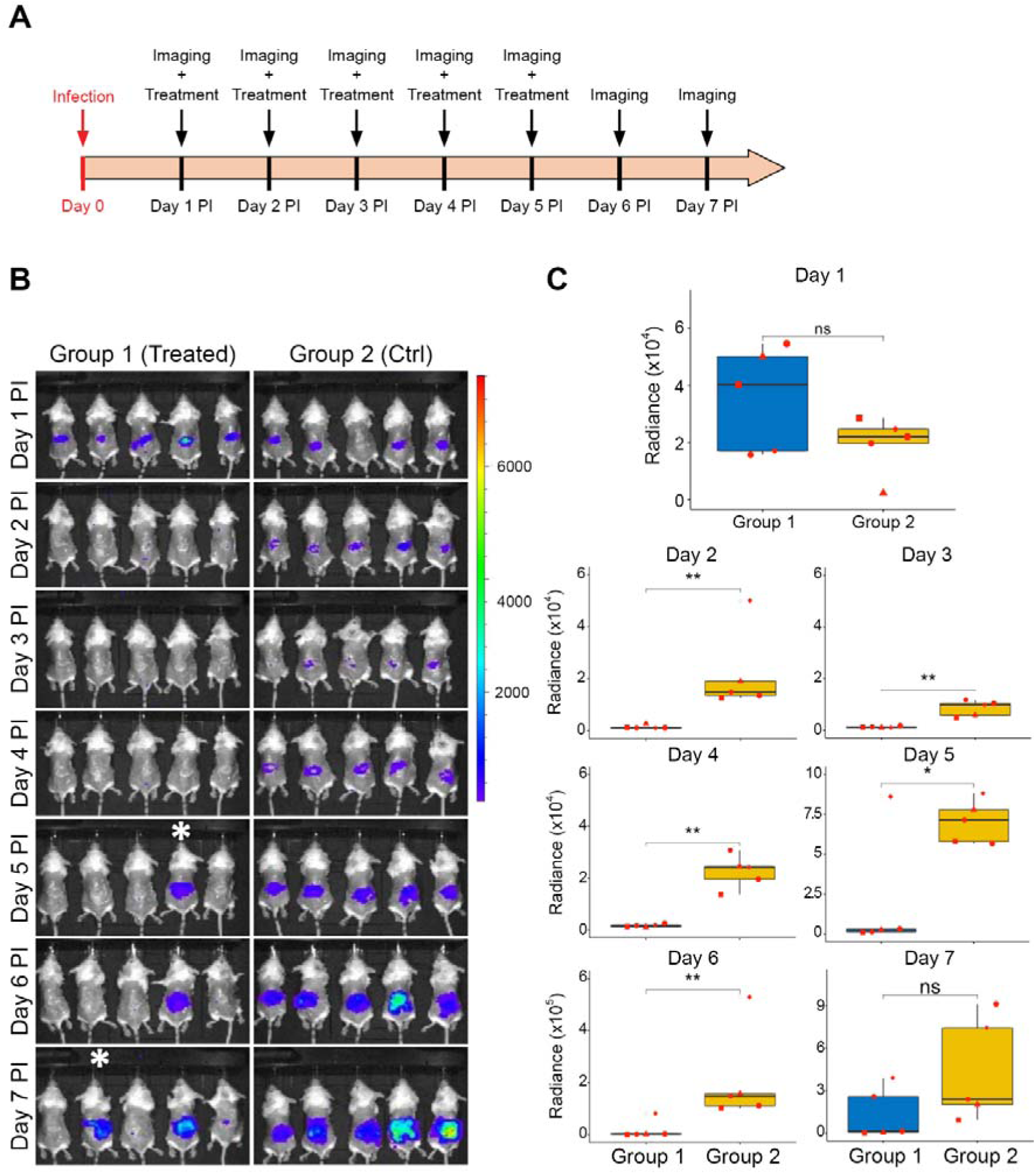
*In vivo* assessment of the efficacy of the combination of micrococcin P1, nisin A and AuresinePlus (MNA) in a skin wound mouse model. The concentration of antimicrobials is the same as D0 in Figure 3B and the treatment was performed from day 1 post-infection (PI) to day 5 PI, imaging and bioluminescence measurements from day 1 was taken prior to initiating treatment. (a) A schematic representation of the experimental setup. (b) *In vivo* imaging of the bioluminescent signals (in photons per square centimeter per steradian) produced by *S. aureus* Xen31 over the 7 day experiment in the treated group 1 and untreated group 2 (Ctrl). A relapse of the infection was seen in two mice of the trated group 1, indicated by white asterisks. (c) quantification of the bioluminescent signals. Statistical significance between experimental groups was assessed with Welch’s t-test. Asterisk representation of statistical significance: *p ≤ 0.05; **p ≤ 0.01; ns = not significant.

Bioluminescence imaging of mice in both groups prior to the initial treatment showed no significant difference between the two groups (Figure 4A, *p* = 0.13). However, following treatment with the MP1/Nis/Aur combination (at concentrations equivalent to D0 in Figure 3) we observed a substantial reduction in bioluminescence in all mice of group 1 (Figure 4B). The bioluminescent signals in the MP1/Nis/Aur-treated group remained significantly lower than mice in control group 2 up to day 6 PI (Figure 4C). However, from day 5 PI and onwards, bioluminescence signals indicated a relapse of the infection in mice of group 1, one mouse by day 5 PI and two mice by day 7, indicating possible resistance development to the antimicrobial combination (Figure 4B, white asterisks).

To further investigate this, bacteria were recovered from the wounds of the two mice (isolates Mut1 and Mut2) and their susceptibility to MP1/Nis/Aur reassessed by a spot-on-lawn assay. As shown in Figure S9, a smaller zone of inhibition was seen for both isolates for the combination (MNA) compared to the wild type strain Xen31. Inhibition zones were particularly reduced for the individual components MP1 and Nis, where one isolate (Mut2) appeared insensitive to MP1 at the concentration tested. Susceptibility to Aur appeared to be similar to the wild type for both isolates. To explain the resistant phenotype of the two isolates, whole genome sequencing was performed to identify potential mutations (Table 1).

**Table 1.** Mutations found in *S. aureus* Xen31-derived isolates showing resistance towards the MP1/Nis/Aur antimicrobial combination in the wound and decreased sensitivity towards MP1 and Nis *in vitro*.

| Strain | Mutation | Consequence on protein | Gene |
| --- | --- | --- | --- |
| Xen31 Mut1 | c.76C>T | p.Pro26Ser | *rplK* |
| Xen31 Mut2 | c.82_93dupTTAGGTCAAGCA | p.A31_G32insLGQA | *rplK* |

c.: chromosome. p.: protein. dup: duplication. ins: insertion.

Both isolates were found to differ from the wild type strain only in the *rplK* gene, encoding the 50S ribosomal protein L11 (RefSeq accession YP_039991.1). Mutant 1 (Mut1) had a missense mutation causing an amino acid change from proline to serine (P26S), while mutant 2 (Mut2) was found to have an in-frame duplication of 12 nucleotides causing an insertion of four amino acids (LeuGlyGlnAla) between amino acids at positions 31 and 32.

### Effect of the combination of micrococcin P1, nisin A and AuresinePlus against *S. aureus* in a murine mastitis model

Mastitis models take advantage of the conserved structure and function of the mammary gland epithelium across mammals and have proven valuable as a tool to investigate host-pathogen interactions within the glands.^58,59^ Conventionally, lactating mice are used for this procedure, and the infection is induced by direct injection of bacteria or other mastitis-causing agents within the mammary gland through the teat canal. In order to develop the mastitis model in our laboratory, lactating CD-1 female mice were subjected to intramammary injection of 10^3^ CFU of *S. aureus* Xen31 at 5-10 days after parturition. The infection was then monitored, with or without treatment, up to 7 days PI. As can be seen in Figure 5A at 1 day PI, both the uninfected and infected mammary tissues show prominent lobular proliferation with dilated lobules containing eosinophilic material consistent with lactational secretions.

**Figure 5.**
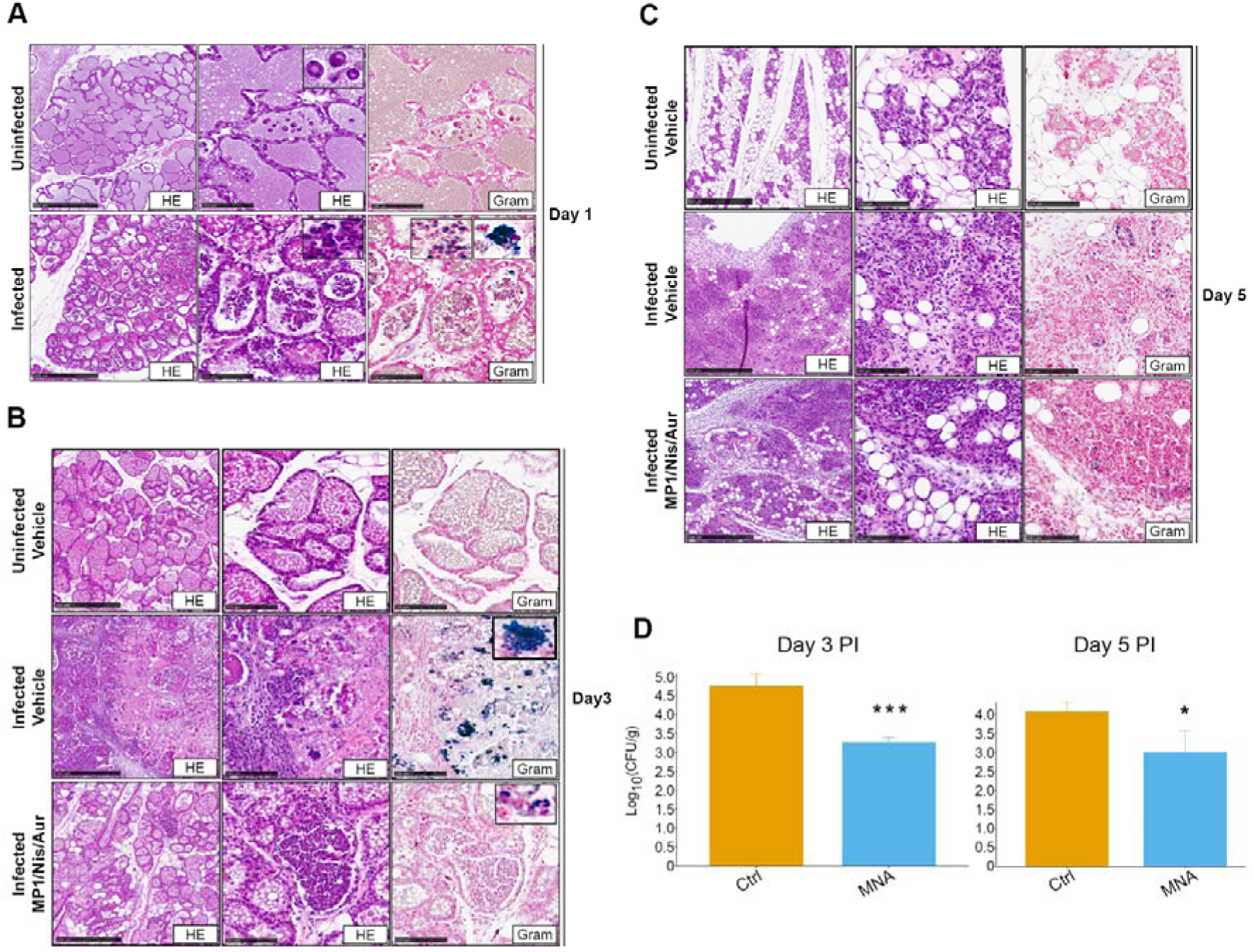
*In vivo* assessment of the efficacy of the combination of micrococcin P1, nisin A and AuresinePlus in a mouse mastitis model. Representative images of the histological examination of explanted mammary glands. (a) from untreated *S. aureus* Xen31-infected and uninfected mice. (b-c) Uninfected mouse injected with the antimicrobial vehicle, and infected mice treated with either the antimicrobial vehicle or the antimicrobial combination at the indicated time-points. Note that the injections of the antimicrobials or their vehicles began at day 1 PI and was extended up to day 5 PI. The concentration used was the same as that corresponding to D0 in Figure 3B. Sequential 5 µm thick tissue sections were stained with either hematoxylin and eosin (HE) or Gram stain (Gram) and images were taken at 5x (left panels) or 20x (middle and right panels). Scale bars represent 500 µm in 5x images and 100 µm in 20x images. Insets show magnified images of areas of interest. (d) Barplots showing the of log10-transformed average colony forming units (Log_10_CFU/g ± 1 standard deviation [error bars]) of bacteria recovered from homogenized mammary glands at day 3 PI (left) and day 5 PI (right). Statistical significance relative to the infected control (vehicle) was assessed with Welch’s t-test. Asterisk representation of statistical significance: *p ≤ 0.05; ***p ≤ 0.001.

The infected mammary tissues show acute immune reaction as evidenced by the presence of neutrophilic infiltrates within the lobular acini as visualized by histochemical staining (hematoxylin and eosin [HE]) (Figure 5A). Gram staining of the same tissues highlighted Gram-positive cells within the neutrophils (Figure 5A, bottom-right panel). These cells appeared to reside predominantly either within phagocytic cells or as larger cell agglomerates resembling biofilm structures.^60^ At 3 days PI, after receiving the first two rounds of treatment (day 1 and 2 PI, see Materials and Methods) the vehicle-treated, uninfected mice (Figure 5B, top panels) showed intact lobules and acini containing lactational secretions. Conversely, glands of vehicle-treated infected mice show necrosis of the acini and the lobular stroma, hemorrhage and marked inflammatory neutrophilic infiltrates of the lobules (Figure 5B, middle panels). Bacterial clumps were present in the lobular acini which resembled biofilm-like structures within the infected tissue. In contrast, the glands of mice treated with the MP1/Nis/Aur combination showed neutrophilic infiltrates limited within the acini and fewer foci of Gram-positive organisms. Furthermore, mice in the treated group had a reduced bacterial CFU count compared to vehicle treated mice (Figure 5B, bottom panels and Figure 5D). By day 5 PI, all tissue samples displayed lobular involution with significant reduction of milk secretion in the acini. Lymphoplasmacytic infiltrates could be observed in the lobules of the infected mice and intralobular fibrosis was more pronounced in the infected vechicle-treated mice than in the infected MP1/Nis/Aur-treated mice (Figure 5C, middle and bottom panels, respectively). A significant reduction in bacterial counts (CFU) from isolated mammary tissues from mice treated with the Nis/MP1/Aur combination was also observed at day 5 PI, although the reduction was less significant compared to day 3 PI (Figure 5D). By day 7 PI, no Gram-positive cells could be detected in vehicle-treated or antimicrobial-treated glands and both displayed a similar morphology to the uninfected vehicle-treated glands (Figure S10A). Treatment of the mammary glands with the MP1/Nis/Aur combination, in the absence of bacterial infection, did not markedly alter the tissue as the appearance compared with the uninfected and untreated controls were similar for the respective time-points (compare Figure S10B).

## Discussion

Mastitis is the most important disease impacting the dairy industry and the welfare of dairy cattle and is frequently caused by Gram-positive bacteria such as *Streptococcus dysgalactiae*, *Streptococcus uberis* and *S. aureus*.^61^ In most cases of mastitis, the disease is sub-clinical and is manifested only with mild symptoms. Nevertheless, in many regions of the world sub-clinical mastitis is treated with antibiotics often without veterinary input.^62^ However, with increasing awareness on antimicrobial resistance, there is a demand for alternative treatment strategies.^61^ In this study, we evaluated an antibiotic-free antimicrobial combination for combating mastitis-associated pathogens both *in vitro* and *in vivo*.

Initially, a combination of bacteriocins micrococcin P1 and nisin was assayed against a panel of mastitis-derived bacteria (Figure 1A). This combination provided good inhibition against most species for a 5 h treatment with low MIC values *in vitro*, except for *S. aureus* which generally exhibited a lower susceptibility to the bacteriocins (higher MIC). However, at 24, 48 and 168 h post-treatment, inhibition was only observed at very high concentrations of the antimicrobials. To extend the effectiveness of the antimicrobial treatment beyond 5 h, the inclusion of enterocin EJ97s (EJs) or garvicin KS was tested. Both tricomponent combinations provided improved inhibition particularly at 48 and 168 h (Figure 2A), but neither combination proved effective against staphylococci and *S. aureus* in particular. To overcome this limitation we included AuresinePlus to the bacteriocin combination. This is an engineered peptidoglycan hydrolase with high potency and specificity against staphylococcal species such as *S. aureus*.^41^ AuresinePlus, together with the MP1/Nis combination provided a significant reduction in MIC against all *S. aureus* isolates at all time-points past 5 h compared with the other combinations (Figure 2A). The reduction in MIC for other staphylococcal species was also observed but the effect was not as pronounced as for *S. aureus*, highlighting the specificity of AuresinePlus (Figure 2B).

The antimicrobial combinations of MP1, Nis and AuresinePlus were additionally shown to be effective at eradicating *S. aureus* biofilms (Figure 3B-C). This led us to assess the antimicrobial efficacy of the combination *in vivo* using a murine skin infection model. For infected mice treated with the combination, a reduction in the bioluminescence signal was observed throughout the experiment, except for two mice where a relapse of the infection occurred (see discussion below, Figure 4C). Thus, our tripartite bacteriocin/PGH combination effectively inhibited the growth of *S. aureus* Xen31 *in vivo*. In the present study we did not correlate bioluminscence with the amount of bacteria present in the wounds. Furthermore, we did not assess possible cytotoxicity of the combination and its effect on wound healing. However, similar studies have been performed on the individual components nisin and MP1, and the enzymes lysostaphin and ClyS which has comparable structure and function to AuresinePlus.^31,34,63–65^ A study by van Staden et al.^66^ reported that skin wounds infected with *S. aureus* Xen36 treated with nisin at a concentration comparable to that used in our study showed reduced bioluminscence and a 2-log reduction in bacterial counts with no negative effects on wound healing. Additionally, MP1 has previously been assessed for topical treatment of skin wounds in mice infected with *S. aureus* Xen31 as part of a formulation (0.1 mg/ml MP1, 5 mg/ml garvicin KS, 5 mg/ml penicillin G in 5% hydropropylcellulose),^31^ which also showed no obvious negative effects on the mice. Another engineered chimeric enzyme ClyS (chimeric lysin for staphylococci) incorporated into a topical ointment was shown to reduce *S. aureus* MW2 by 3-log (single dose of 10% wt/wt) in a murine skin infection model, with no macroscopic alterations to the wound.^63^

A tripartite bacteriocin/PGH combination in which each of the components have distinct mechanism of action, such as the ones used here, will not only be more effective due to its expanded antimicrobial spectrum, but it will also help counteract potential shortcomings of the individiual components such as instability and emergence of mutants. As individual components, these antimicrobials probably have limited efficacy against *S. aureus* in wounds. AuresinePlus is an engineered PGH consisting of the catalytic domain of the S. aureus autolysin LytM and the SH3b cell-wall binding domain lysostaphin from *S. simulans*.^48^ Lysostaphin belongs to a class of large (> 10 kDa) heat-labile bacteriolysins, consequently, these enzymes are sensitive towards proteases. It is well-known that wounds are protease rich environments, the lower efficacy of AuresinePlus in the wound could therefore be due to degradation of AuresinePlus in the wound.^67^ Furthermore, spontaneous mutations conferring high levels of nisin tolerance to *S. aureus* has been reported to occur at a frequency of 2×10^-7^ *in vitro*.^68^ Additionally, spontaneous mutants resistant to MP1 are frequently found *in vitro*, but can also arise *in vivo*.^31,69,70^ Thus, while *S. aureus* can resist treatment with the individual components, it is highly improbable that the same *S. aureus* cells can evade all three components simultaneously.

Nevertheless, as mentioned above, two of the infected mice treated with the MP1/Nis/Aur combination showed an increase in bioluminescence after day 5 PI, suggesting a relapse of the infection. Suspecting that cells in these mice had developed resistance to one or more of the components in the treatment, whole-genome sequencing was performed on an isolate from each mouse. Both isolates harbored mutations in *rplK*, encoding the ribosomal protein L11. This was not unexpected, as the molecular target of MP1 is the *rplK*-encoded ribosomal protein L11, and certain mutations in this gene are known to confer resistance to MP1 and other thiopeptides.^71^ Binding of MP1 interferes with the interaction between ribosomal proteins L11 and L7 which inhibits a translocation step induced by hydrolysis of elongation factor G.^71^ Both mutants retained the sensitivity to AuresinePlus. Notably, however, the two isolated *rplK* mutants showed cross-resistance also to nisin. To our knowledge, mutations in *rplK* have not previously been shown to reduce susceptibility towards nisin. Nisin inhibits growth of bacteria by binding to and sequestering lipid II, an essential precursor molecule for cell wall biosynthesis.^72^ Additionally, at higher concentrations, nisin in complex with lipid II can oligomerize into a pore-forming complex resulting in a loss of membrane potential and leakage of intracellular contents.^73^ It is uncertain how mutations in *rplK* confers tolerance towards nisin. It could be speculated that reduced growth rate in these mutants resulting from lower translation efficiency can partially compensate for the inhibition of cell wall synthesis and lead to persistence. Mutations associated with nisin resistance has previously shown to occur in genes encoding the two-component stress response system BraRS or the response regulator PmtR.^74–76^ It is not known if mutations in BraRS affects the fitness or virulence of *S. aureus*, but a mutant of *pmtR* showed reduced virulence and survivability in a mouse infection model, effectively limiting the problem with the emergence of such mutations *in vivo*.^77^

Encouraged by the efficacy of the MP1/Nis/Aur combination in inhibiting *S. aureus* in the wound, we sought to examine the potential of the antimicrobial combination in treating mastitis, an infection often caused by *S. aureus*. Intramammary injection of the combination significantly reduced bacterial counts compared to untreated controls, clearly demonstrating the effectiveness of the tripartite combination in reducing the intramammary infection (Figure 4C). Furthermore, intramammary injection of the combination in uninfected mice showed no considerable morphologic changes in the glandular architecture of the breast tissues (Figure 5), suggesting low cytotoxicity of the combination at the concentration tested. Lysostaphin has been documented to be well-tolerated by normal human epidermal keratinocytes with IC_50_ (midpoint cytotoxicity value) of 16 mg/ml.^78^ However, nisin and micrococcin P1 exhibit some cytotoxicity to eukaryotic cell lines in *in vitro* assays with IC_50_ of 0.3-0.4 mg/ml for nisin (HT29 and Caco-2 cell lines),^79^ and micrococcin P1 impairs growth of HepG2 and THP-1 cell lines at concentrations above 0.03 mg/ml.^28^ Potential negative effects of the combination on the mammary tissue is likely reduced by the immediate diffusion and dilution of the components within the cells and interstitial fluid. However, we cannot exclude cytotoxic effects that are not easily apparent from histologic examination with H&E stain.

Taken together, these assays demonstrate that the bacteriocin/PGH combination can effectively treat infections *in vivo*. However, for an optimal and long-lasting effect, proper antimicrobial dose and administration frequency are critical. We expect antimicrobial combinations to play an increasingly important role in combating multidrug-resistant bacteria. In contrast to monotherapeutic strategies, combinations of several antimicrobials have the potential of broader inhibition spectra and to reduce chances of resistance development. However, the development of resistance to alternative antimicrobials such as bacteriocins and emergence of more general cross-resistance mechanism will require more research. By employing antimicrobials with different non-overlapping mechanisms of action, resistance development becomes exceedingly unlikely. As with other drugs, further development of the treatment tested here will also require more knowledge about the cytotoxicity and the pharmacodynamics of the antimicrobial combination. Furthermore, by utilizing antibiotic-free antimicrobial agents when possible, we may hope to prolong the effectiveness of current antibiotics.

## Materials and methods

### Bacterial Strains

All bovine mastitis-derived strains have been obtained from Tine (Molde, Norway) with the exception of *S. aureus* MOK strains and RF 122 which were a kind gift from Orla Keane (Teagasc, Oak Park, Carlow, Ireland). All strains were routinely grown in brain heart infusion (BHI) broth (Oxoid, United Kingdom) at 37 °C under aerobic conditions without shaking, with the exception of *T. pyogenes* strains which were grown in BHI supplemented with 5% heat-inactivated fetal bovine serum (VWR). For *in vivo* imaging of bacterial infection in mice, *S. aureus* Xen31 (Perkin Elmer, Waltham, MA) was used. The strain was derived from the parental strain *S. aureus* ATCC 33591, a clinical MRSA strain isolated from Elmhurst Hospital in New York, NY, USA.^80^ *S. aureus* Xen31 possesses a stable copy of the modified *Photorhabdus luminescens luxABCDE* operon at a single integration site on the bacterial chromosome.

### Antimicrobials

GarKS peptides and EJs was synthesized by Pepmic Co., Ltd. (China) with ≥95% purity and solubilized to concentrations of 1□mg/ml in sterile Milli-Q water. Micrococcin P1 was produced as described previously and solubilized to a concentration of 10 mg/ml in dimethyl sulfoxide (Sigma-Aldrich).^30^ Nisin A was obtained as part of a commercial fermentate which was solubilized to a concentration of 1 mg/ml in a 0.05% acetic acid solution (Merck). AuresinePlus was solubilized to a stock concentration 1 mg/ml in a buffer containing 20 mM Tris-HCl pH 7.0, 200 mM NaCl, 10% glycerol. All antimicrobials were diluted to their working concentrations on the day of the experiments, stock solutions were stored at −20 °C until use. Hydroxypropyl cellulose (HPC) with a weight-average molecular weight (*M*w) of □80,000 g/mol and a number-average molecular weight (*M*n) of □10,000 g/mol was used (Merck).

### Planktonic cell growth inhibition assays

Growth inhibition assays were performed in 96-well microtiter plates (Sarstedt). Briefly, 135□μl of growth medium were dispensed in each well of the plate (according to the number of bacterial strains tested) except in the wells of the first row. The antimicrobials were diluted in the growth medium to working concentrations (see below) in a final volume of 285□μl and dispensed in the wells of the first row. From the first row, 150□μl of the antimicrobials were then serially diluted (twofold dilutions) in a sequential fashion until the last row of the plate. Finally, 15□μl of a fresh O/N culture of each strain were added in the appropriate wells to reach a final volume of 150□μl in each well. The plates were then incubated at 37□°C for 5, 24, 48 or 168□h. The growth inhibition was expressed as a minimum inhibitory concentration (MIC), which refers to the minimum concentration of the antimicrobial needed to abolish the bacterial cell growth. The MIC values were assessed by optical density readings at 600□nm (O.D.600).

The growth media used for the assays were BHI for all the strains with the exception of *T. pyogenes* strains (see above). The working concentrations of the antimicrobials were as follows: 100□μg/ml for garvicin KS, EJs, nisin A and AuresinePlus and 10 μg/ml for micrococcin P1.

### *In vitro* biofilm production and biofilm formation ability assay

Ten microlitres of the O/N cultures were diluted 1/10 in 90□μl of the appriopriate medium and and dispensed in the wells of a 96-well microtiter plate (Sarstedt) to a final volume of 100□μl. The plates were then incubated at 37□°C for 24□h. After the incubation, the pres-ence of the biofilm at the bottom of the wells was initially confirmed visually. Biofilm for-mation ability assays were performed as described previously.^32,34^ Bacterial biofilms were allowed to form for 24□h prior to being washed twice with 100□μl of 0.9% NaCl at room temperature (RT) to remove planktonic cells. Biofilms were then left to air dry for 15□min. After drying, 200□μl of a 0.4% solution of crystal violet (Merck) were added to each well and incubated for an additional 15□min. The dye was then removed and the wells were washed three times with 200□μl of 0.9% NaCl, bound crystal violet was then extracted by incubating the wells with 100□μl 70% ethanol. The extraction procedure was repeated twice, and the combined crystal violet amount extracted was quantified by optical density measurements at 600□nm. The quantification of the crystal violet released from the biofilm is a surrogate measure of the number of bacterial cells forming the biofilm.

Media used for the biofilm formation were as follows: tryptic soy broth (TSB – Sigma Aldrich) supplemented with 1% glucose and 1% NaCl for all *S. aureus* strains; TSB supplemented with 1% glucose for all other for all other species with the exception of *T. pyogenes* strains, for which BHI supplemented with 5% fetal bovine serum was used.

### Biofilm-oriented antimicrobial test (BOAT)

The BOAT assays were essentially performed as described previously.^32,34^ Serial dilutions of the antimicrobials were prepared in challenge plates as follows: 175□μl of TSB were trans-ferred in each row of a 96-well microtiter plate, except for the first row, according to the number of microbial strains tested. In the first row of the plate, the antimicrobials were dilut-ed to their respective working concentrations in a final volume of 350□μl of TSB. From the first row, 175□μl of the antimicrobial dilutions were then transferred to the second row of the plate and further serially diluted to the bottom of the plate. The same procedure was followed to prepare the controls, except that instead of the antimicrobials an equivalent volume of the respective vehicles was used. Unless otherwise stated, the starting concentrations of the antimicrobials for all experiments involving biofilms were 1□mg/ml nisin A, 0.1□mg/ml for micrococcin P1 and AuresinePlus and 0.65□mg/ml for EJs.

Bacterial biofilms were produced as described above. After washing the biofilm twice with 100□μl of sterile saline, a total of 150□μl of the antimicrobial and control dilutions were transferred from the challenge plate to the corresponding wells of the biofilm plate. The chal-lenged biofilms were then incubated for an additional 24□h at 37□°C. After the challenge period, the antimicrobial dilutions were removed and the biofilms were carefully washed three times with 150□μl of the sterile saline. A total of 100□μl of TSB supplemented with 0.025% of triphenyl-tetrazolium chloride (TTC, Sigma-Aldrich) were then added to each well of the plate and further incubated at 37□°C for 5□h. The results were then assessed by monitoring the development (or not) of formazan (red color), a measure of metabolic activity by the bacterial cells. The medium was then removed and the red formazan was solubilized by adding 200□μl of ethanol:acetone (70:30) mixture to each well and incubated O/N. The amount of extracted dye, reflecting the degree of bacterial cell metabolic activity, was then quantified by spectrophotometric readings at 492□nm.

### Determination of the bacterial viability after BOAT

The procedure for the BOAT assay was repeated as described above except that instead of adding the TTC solution, the antimicrobial-challenged cells were resuspended in TSB and then serially diluted in TSB buffer. Serial dilutions of the bacterial cells were then plated on BHI agar plates and incubated at 37□°C for 24□h. The results were then assessed by direct counting of the developed colonies and the CFU was determined.

### Mice

Animal experiments were approved by the Norwegian Food Safety Authority (Oslo, Norway FOTS ID 29800) and the experiment was executed in accordance with the Norwegian Regulation concerning the use of animals for scientific purposes and EU Directive 2010/63.47. Two to four mice were housed per cage in individually ventilated cages (IVCs; Innovive Inc., San Diego, CA) during the whole experiment and maintained on a 12 h light/12 h dark cycle with *ad libitum* access to water and a regular chow diet (RM1; SDS Diet, Essex, United Kingdom). Cages were equipped with chewing sticks and igloo houses with a running wheel, and kept at a temperature of 25±1 °C and relative humidity of 50±5%. The mice were acclimatized in our mouse facilities for at least 1 week before the start of experiments.

### Mouse skin infection model

Before infection and treatment, the mice were anesthetized and shaved on the back and flanks. Anesthetics was an injectable cocktail containing Zoletil Forte (Virbac, Carros, France), Rompun (Bayer, Oslo, Norway), and Fentadon (Eurovet Animal Health, Bladel, The Netherlands) cocktail (ZRF) (containing 3.3□mg Zoletyl Forte, 0.5□mg Rompun, and 2.6□μg fentanyl per ml 0.9% NaCl) by intraperitoneal injection (0.1□ml ZRF/10 g body weight) and shaved on the back and flanks with an electric razor. The remaining hair was removed with hair removal cream (Veet; Reckitt Benckiser, Slough, United Kingdom) according to the manufacturer’s instructions. The next day, the mice were again anesthetized with ZRF cock-tail (0.1□ml/10 g body weight), and two skin wounds were made on the back of each mouse with a sterile biopsy punch 6□mm in diameter (dermal biopsy punch; Miltex Inc., Bethpage, NY). Prior to infection, overnight-grown *S. aureus* Xen31 cells were washed twice in sterile saline and then suspended in ice-cold phosphate-buffered saline (PBS) buffer. Each wound was inoculated with 10□μl of PBS containing 2×10^7^ CFU of *S. aureus* Xen31 cells using a pipette tip. After bacterial application, the mice were kept on a warm pad for 10 to 15□min to dry the inoculum, and the wounds were then covered with a 40- by□50-mm Tegaderm film (3M Medical Products, St. Paul, MN, USA). The mice were then left for 24 h for the infection to establish. The day after (day 1 post-infection [PI]), the mice were anesthetized with 2% isoflurane, and the luminescent signal was measured with an IVIS Lumina II (Perkin Elmer; 2-min exposure time). The luminescent signal was quantified with Living Image software (Perkin Elmer) from regions of interest (ROI) around the wound and expressed as photons per second per square centimeter per steradian.

Mice were then split into two groups: group 1, mice were treated with the antimicrobial com-bination (1□mg/ml nisin A, 0.1□mg/ml MP1, 0.1□mg/ml AuresinePlus) in a 5% (wt/vol) HPC hydrogel; group 2, mice were treated with the antimicrobial vehicles diluted in 5% (wt/vol) HPC a negative control. Mice were treated daily by injecting the antimicrobials or the vehicle controls (50 µl) directly on the infected wounds through the Tegaderm film. The treatments were administered from day 1 to day 5 PI, whereas the bioluminescence imaging was per-formed daily for the whole course of the experiments (day 1-7 PI).

### Mouse mastitis model

Time-mated pregnant female mice, guaranteed at E15 on delivery date, were purchased from Janvier Labs (France). Four to six lactating females, 5-10 days after parturition, were anesthetized with a constant flow of 2% isoflurane. The abdominal area of the mice was disinfected with a 70% (v/v) ethanol solution and the 5^th^ pair of mammary glands were injected with either 50 µl of a *S. aureus* Xen31 suspension containing 10^3^ CFU in endotoxin-free PBS (VWR), or endotoxin-free PBS alone using a 50 µl Hamilton Neuros 700 Series Microliter Syringe (Hamilton) equipped with a removable 33 G blunt-end needle. Twenty-four hours after injection, mice were divided into four groups: 1, bacterially injected and treated with antimicrobial combination; 2, bacterially injected and treated with vehicle-control; 3, PBS-injected and treated with antimicrobial combination; 4, PBS-injected and treated with vehicle-control. Treatments (antimicrobial combination or vehicle control) were administered once daily from day 1 to day 5 PI via intramammary injection. Starting at day 3 PI and every other day onwards (days 3, 5 and 7) 4 mice per group were sacrificed by cervical dislocation and subjected to surgical removal of the 5^th^ pair of mammary glands. At the same time 1 uninjected lactating female was also sacrificed at the same time-points to serve as a tissue morphology and gland involution control. For each mouse one of the glands were placed in sterile, endotoxin-free PBS and one in a 4% solution of paraformaldehyde (PFA – Sigma Aldrich). Mammary glands stored in PBS were immediately subjected to tissue homogenization using a GentleMACS Dissociator equipped with GentleMACS M tubes (Myltenyi Biotech) following the manufactureŕs instructions. The resulting homogenates were serially diluted in BHI medium and plated on selective BHI-agar plates containing 50 µg/ml ampicillin (Merck). The plates were incubated overnight at 37 °C to allow the growth of ampicillin-resistant colonies that were used for CFU determination. The glands in PFA were stored at 4 °C overnight to allow full tissue fixation followed by histological processing.

### Histology

After fixation in PFA, the tissues were embedded in paraffin and 5 µm sections was obtained using a microtome. Sections were stained with hematoxylin and eosin (H&E) and high resolution images were acquired using an automated slide scanner.

### Sequencing and sequence analysis

Sequencing and library preparation was performed by Novogene (Beijing, China) in paired-end mode (2×150 bp) to an average sequencing depth of approximately 600 for each sample. Sequencing reads obtained from *S. aureus* Xen31 was assembled using Unicycler v0.5.0 with the SPAdes genome assembler v3.15.5.^81,82^ Assembled contigs were annotated using prokka v1.14.6 with the provided database for genus *Staphylococcus*.^83^ Sequencing reads obtained from *S. aureus* Xen31 Mut1 and Mut2 were directly mapped against the annotated contigs for detection of variants using snippy v4.6.0.^84^

### Statistical analysis and data representation

The statistical analysis and graphical representations for all data were performed with R Studio (Version 2023.06.0+421) and R (version 4.3.2) softwares.

## Acknowledgements

We are grateful to Prof. Dzung B. Diep, who passed away December 2022, for initiating this project. We would like to thank Orla Keane (Teagasc, Ireland) and the TINE Mastitis Laboratory (TINE SA, Molde, Norway) for giving access to mastitis-derived strains.

## Funding

This work was funded by a EEA/Norway grant (grant NOR/POLNOR/PrevEco/0021/2019).

## Transparency declarations

The authors declare the following financial interests which may be considered as a potential competing interest: IS is co-founder of Enzybiotx, a company which develops enzyme-based antimicrobials.

The remaining authors declare that the research was conducted in the absence of any commercial or financial relationships that could be construed as a potential conflict of interest.

