## Supplementary materials and methods for "An antibiotic-free antimicrobial combination of bacteriocins and a peptidoglycan hydrolase: *in vitro* and *in vivo* assessment of its efficacy"

**Supplementary Figures**

Kranjec et al.

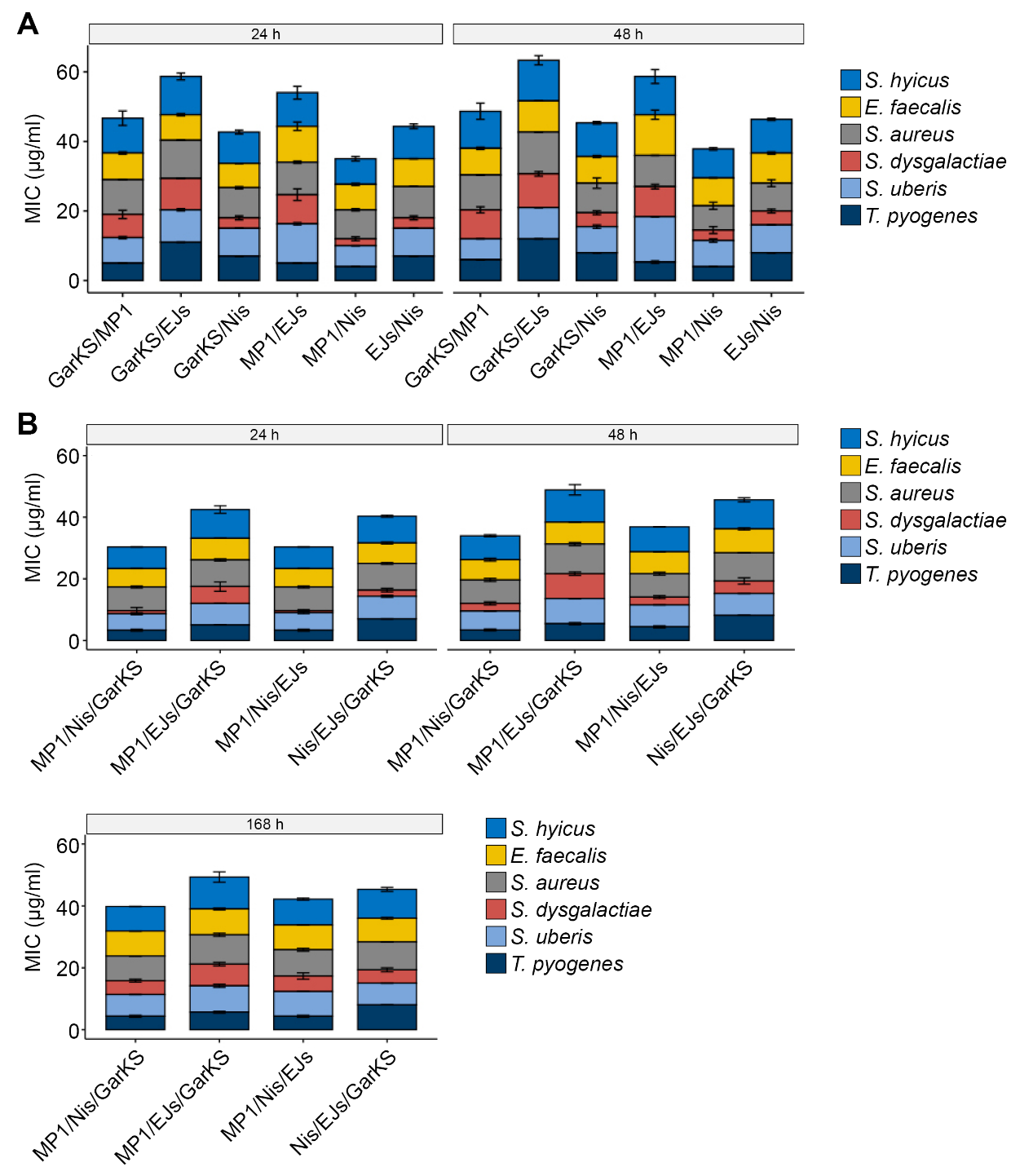

**Figure S1**. Testing of bacteriocin combinations against a panel of mastitis-derived bacterial species. Stacked barplots showing the cumulative MICs of the indicated species against (a) bicomponent, or (b) tricomponent combinations for the indicated time-points.

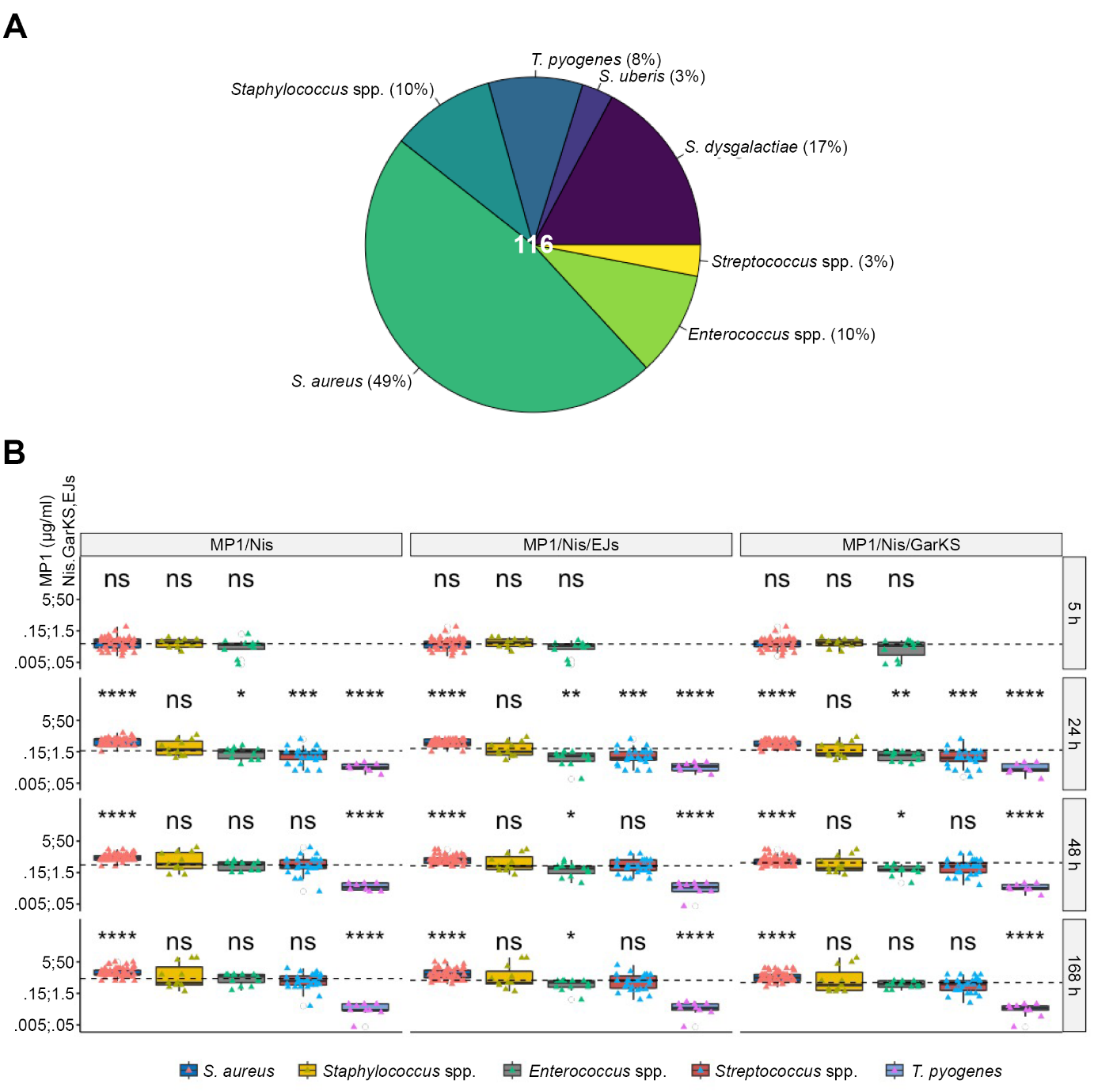

**Figure S2**. Mastitis-derived bacterial collection and susceptibilities to bacteriocin combinations. (a) Pie chart showing the composition of the mastitis bacterial collection used in this study. The species in the collection are indicated with the relative contribution in percentage. (b) Boxplots showing the median planktonic MIC value distributions (thick line within boxes) of the strains belonging to the species indicated in the legend (bottom) and included in the mastitis collection against the bacteriocin combinations indicated on the top. Antimicrobial assays were performed at 5, 24, 48 and 168 hours, as indicated on the right-hand side of the plot. The dotted lines in each plot represent the average MIC values obtained across species for each treatment and for each time-point. For each plot the statistical significance was assessed as deviation from the average MIC value using the Welch’s t-test. Data for *T. pyogenes* at 5 hours are missing due to the lack of measurable bacterial growth of the bacterial strains. Asterisk representation of statistical significance: *p ≤ 0.05; **p ≤ 0.01; ***p ≤ 0.001; **** p ≤ 0.0001; ns = not significant.

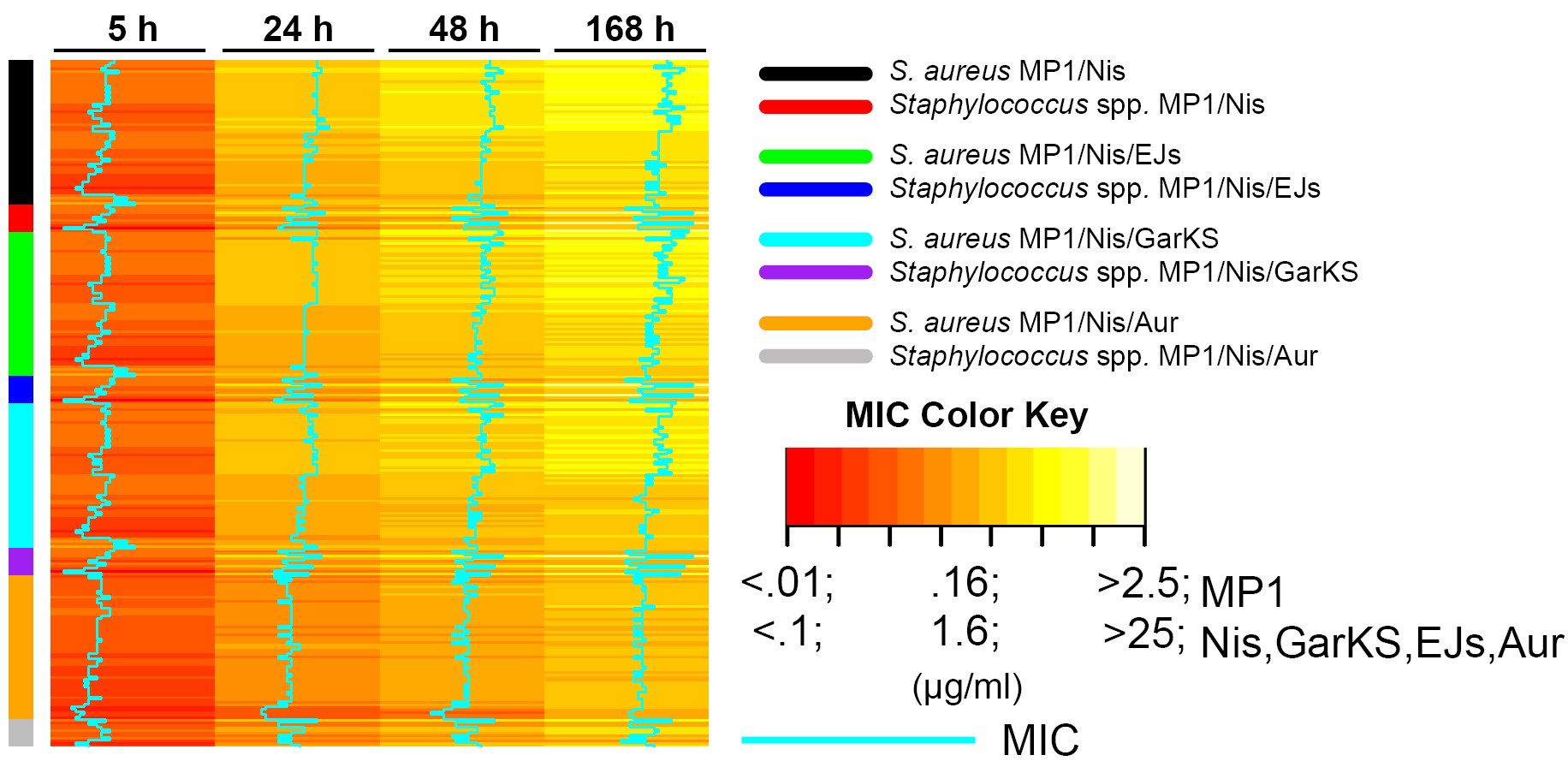

**Figure S3**. Contribution of AuresinePlus to the antimicrobial activity of micrococcin P1 and nisin A against staphylococci. Heatmap showing the distribution of MIC values (µg/ml) of *S. aureus* and other staphylococcal species (spp.) against the indicated antimicrobial combinations.

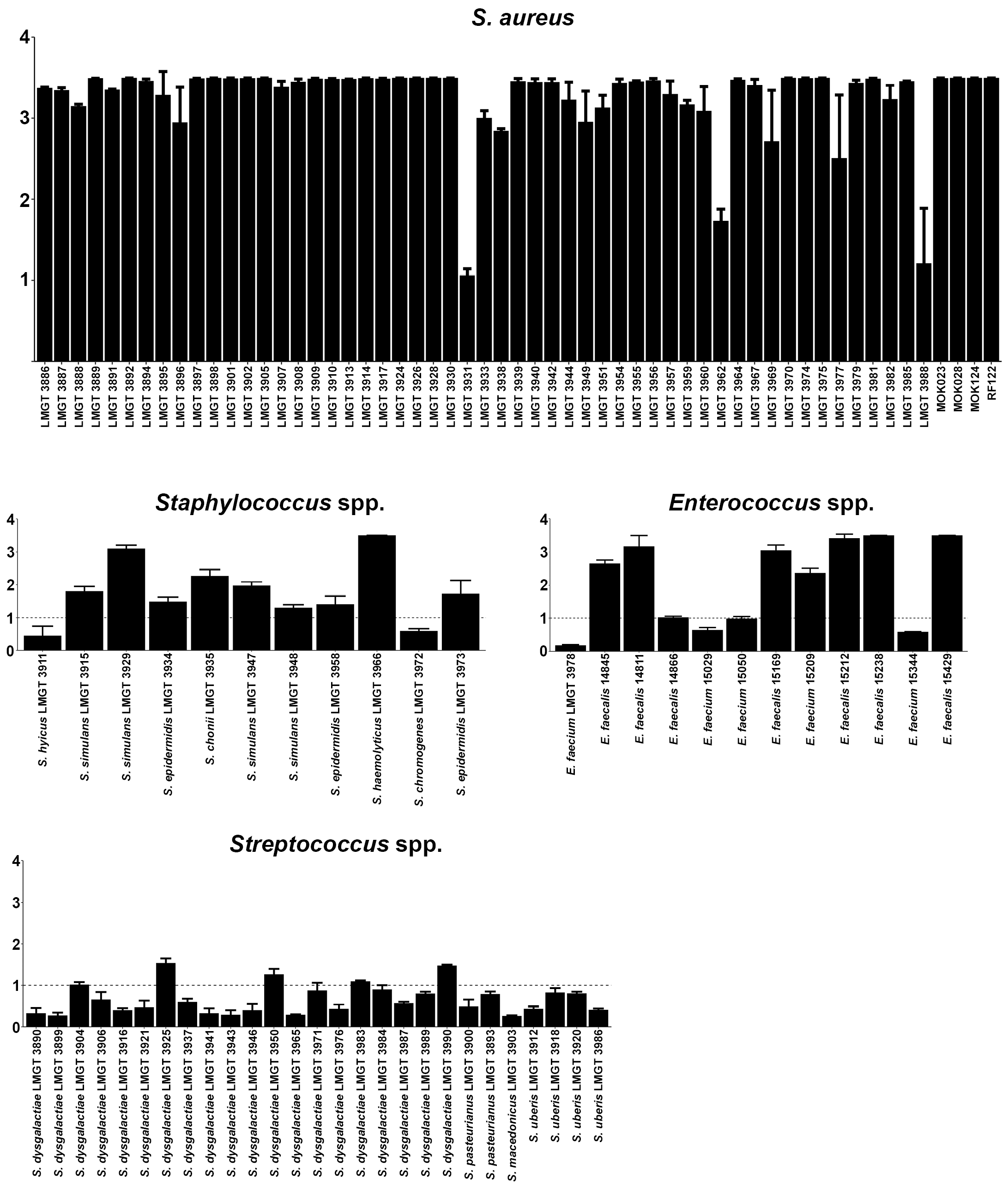

**Figure S4**. Biofilm formation abilities of isolates included in the mastitis collection. Indicated strains were allowed to grow for 24 h in the wells of a 96-well plate prior to crystal violet staining of the formed biofilm. The amount of dye bound to each well was used as an indirect measure of biofilm formation ability and was quantified by optical density readings at 600 nm (OD_600_) for each strain. The average values (± s.d.) are shown. The dotted line corresponds to the OD level below which strains were considered as nonbiofilm formers. Strains were considered moderate, good or strong biofilm formers with OD values over 1, 2 or 3, respectively.

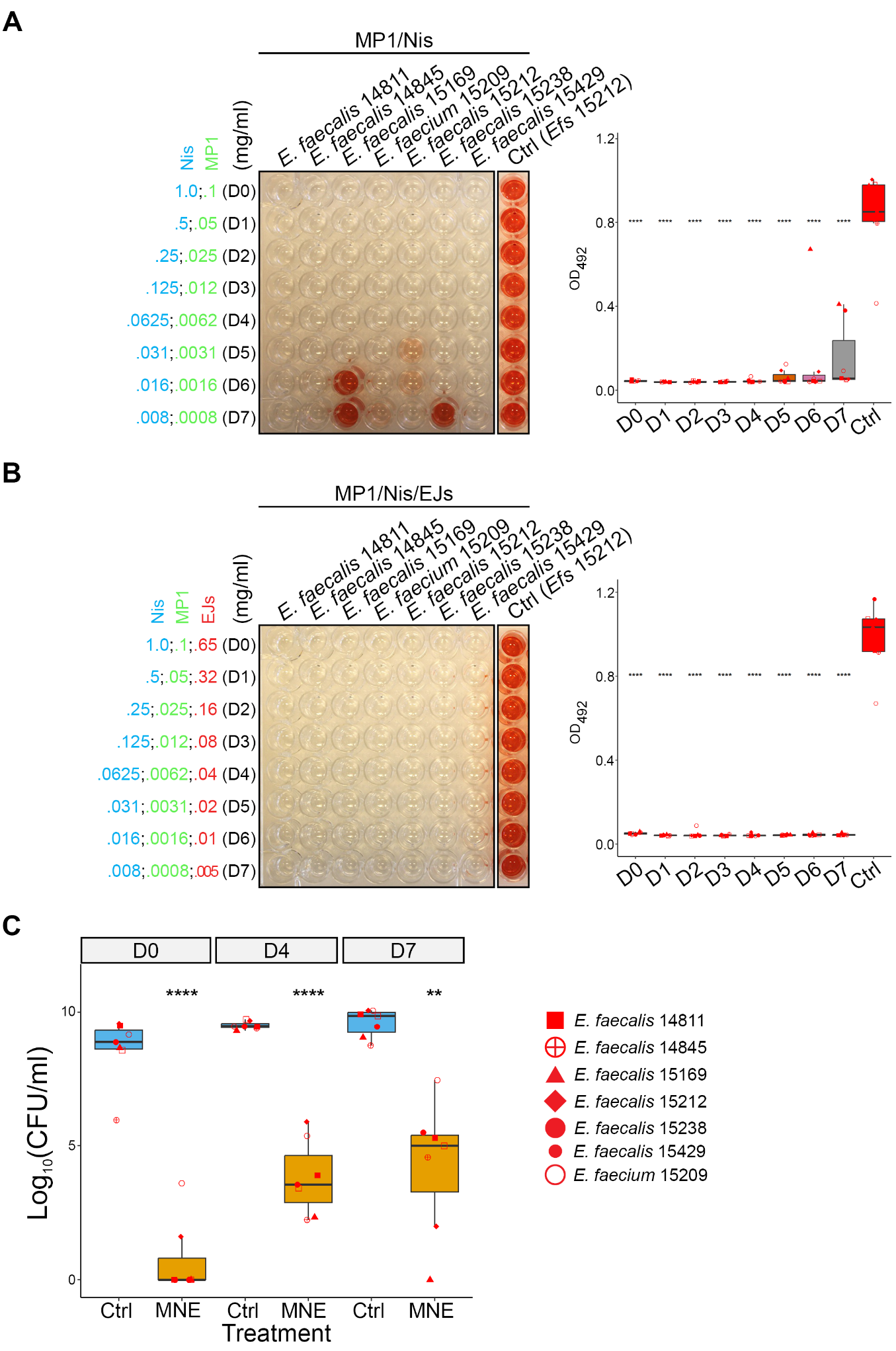

**Figure S5**. Assessment of bacteriocin combinations against mastitis derived enterococci. (a) The left panel shows a representative image of the BOAT assay performed with a twofold dilution of the MP1/Nis combination against the indicated enterococcal strains. The concentrations of the antimicrobials (in mg/ml) are shown on the left (dilutions D0-D7). The assays were simultaneously performed with the control vehicles of the antimicrobials (Ctrl), and a representative image of *E. faecalis* strain 15212 is shown. Red color indicates metabolic activity and its quantification was performed by optical density readings at 492 nm (OD_492_). The box plot (Tukey’s) on the right panel displays the quantification of the recovery of bacterial metabolic activity as a function of the dilution. (b) Same as in panel A but the strains were tested with the MP1/Nis/EJs combination. Post-hoc statistical analyses of each group (D0-D7) relative to the vehicle control (Ctrl) was performed with Welch’s t-test. (c) Boxplot showing the log_10_-transformed colony forming units (Log_10_CFU/ml) calculated after the BOAT assay for the same strains as in panels A and B. The CFU quantification was performed for the MP1/Nis/EJs combination for dilutions D0, D4 and D7 depicted in panel B. Asterisk representation of statistical significance: **p ≤ 0.01; **** p ≤ 0.0001.

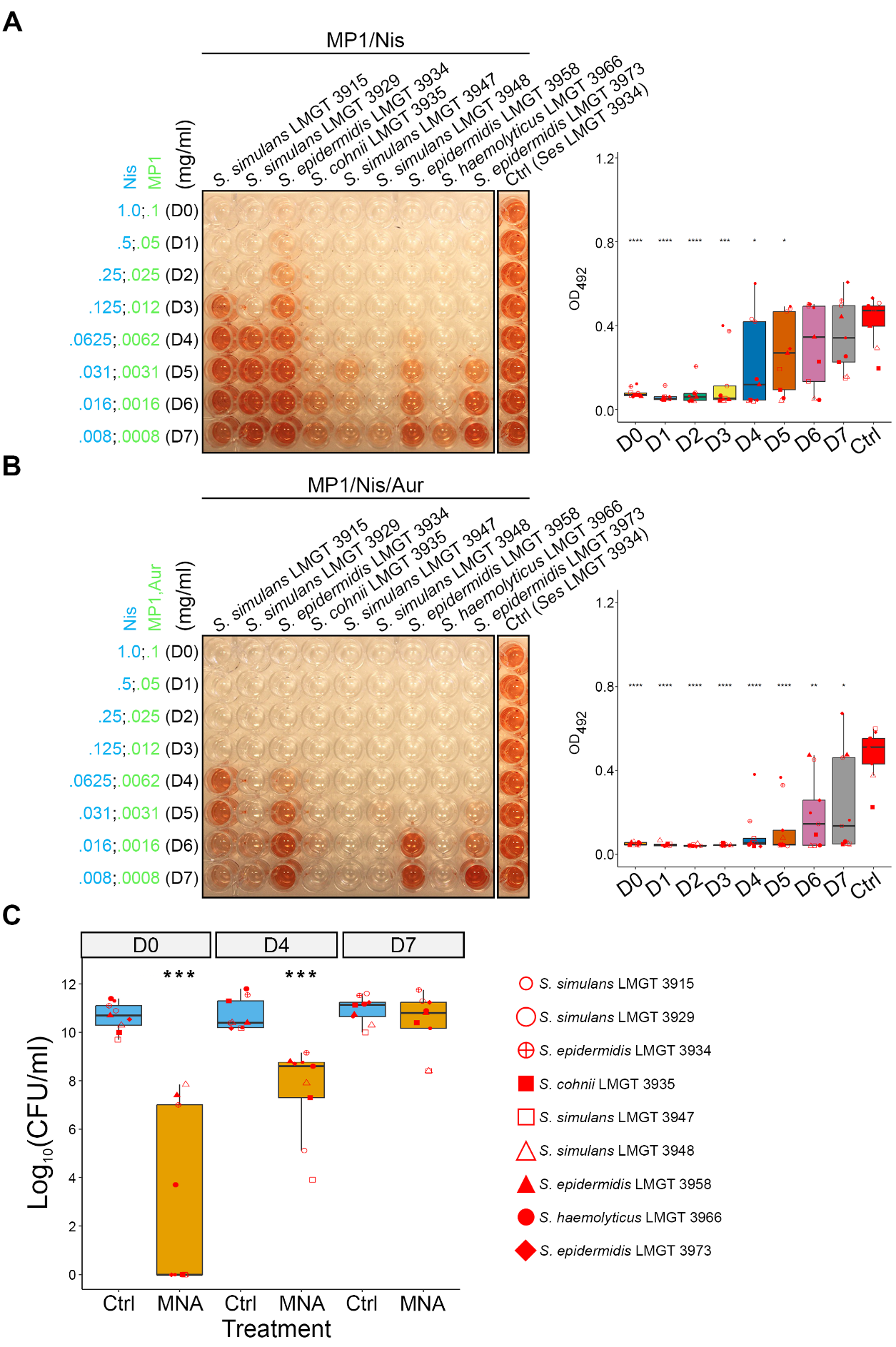

**Figure S6**. Assessment of bacteriocin and bacteriocin-AuresinePlus combinations against of mastitis-derived Staphylococcal species. (a) The left panel shows a representative image of the BOAT assay performed with a twofold dilution of the MP1/Nis combination against the indicated strains. The concentrations of the antimicrobials (in mg/ml) are shown on the left (dilution factors D0-D7). The assays were simultaneously performed with the control vehicles of the antimicrobials (Ctrl), and a representative image of *S. epidermidis* strain 3934 is shown. The development of the red color indicates metabolic activity, and its quantification was performed by optical density readings at 492 nm (OD_492_). The boxplot in the right-hand panel displays the quantification of the recovery of bacterial metabolic activity as a function of the dilution factor. (b) Same as in panel A but the strains were tested against the MP1/Nis/Aur tricomponent combination. The global statistical significance in the plots in panels A and B was assessed using the one-way ANOVA test, whereas the post-hoc statistical analyses of each group (D0-D7) relative to the vehicle control (Ctrl) was performed with the Welch’s t-test. C) Boxplot showing the median distribution of log_10_-transformed colony forming units (Log_10_CFU/ml) calculated after the BOAT assay for the same strains as in panels A and B. The CFU quantification was performed for the MP1/Nis/Aur combination at concentrations D0, D4 and D7 depicted in panel B. Asterisk representation of statistical significance: *p ≤ 0.05; **p ≤ 0.01; ***p ≤ 0.001; **** p ≤ 0.0001.

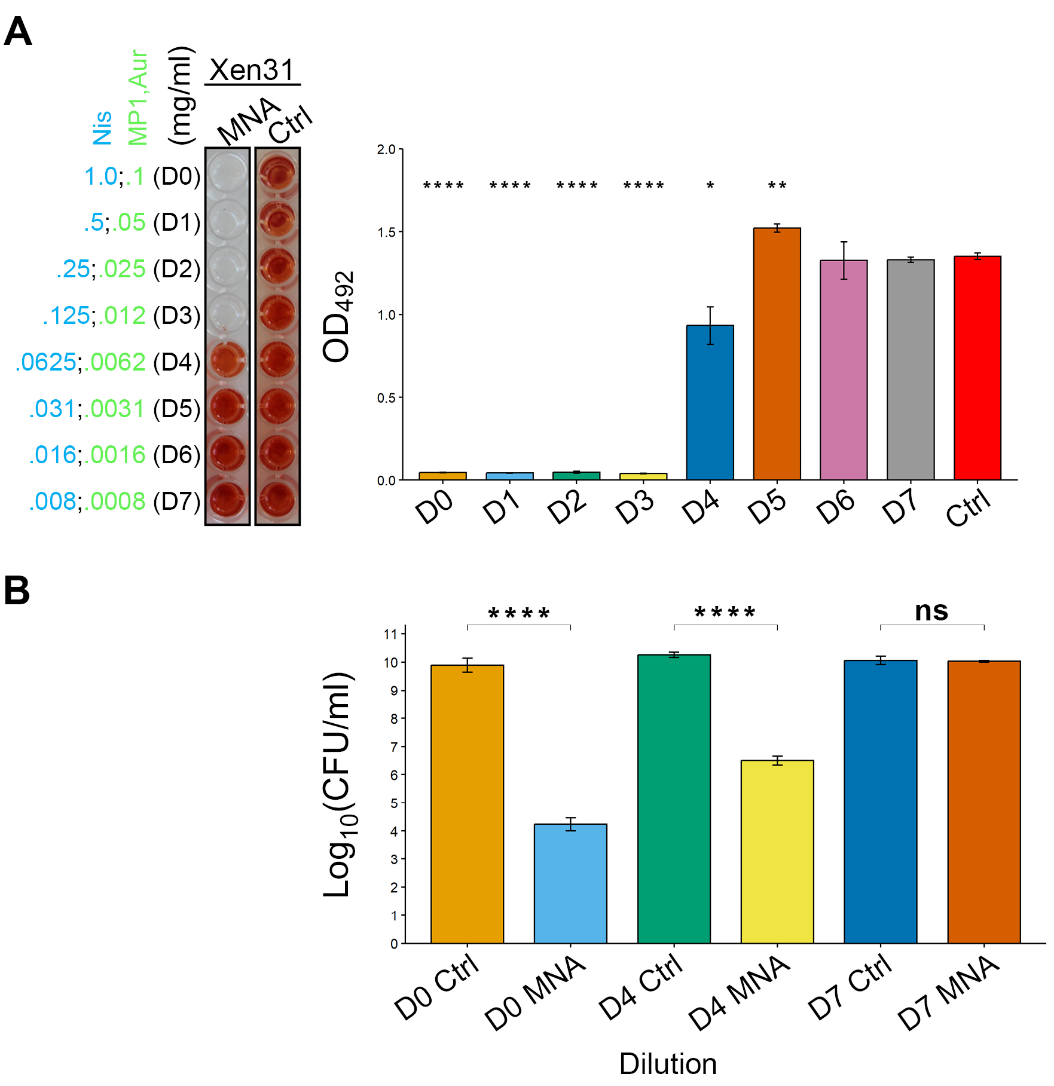

**Figure S7**. Efficacy of the micrococcin P1, nisin A, AuresinePlus combination against *S. aureus* Xen31. (a) Representative image of BOAT assays performed with the antimicrobial combination against the Xen31 strain. The concentrations were the same as in Figure 3 and Figure S6B. The bar plot on the right represents the trend of metabolic activity recovery as a function of the dilution as described in Figures 3 and S6. (b) barplot showing the median distribution of log_10_-transformed colony forming units (Log_10_CFU) values calculated after the BOAT assays in panel A. Asterisk representation of statistical significance: *p ≤ 0.05; **p ≤ 0.01; **** p ≤ 0.0001; ns = not significant.

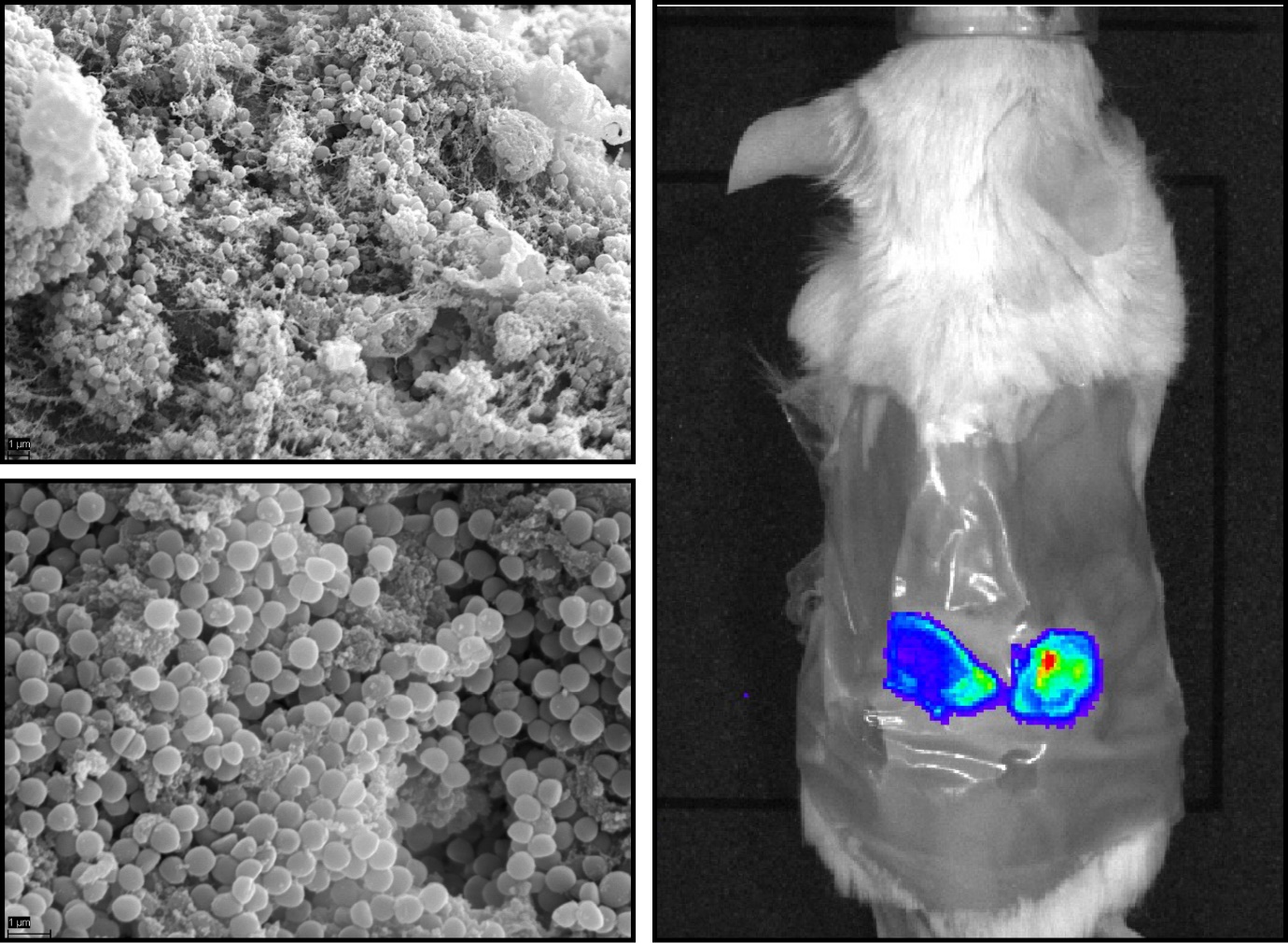

**Figure S8**. Establishment of biofilm associated infection by *S. aureus* Xen31 in the mouse wound model. Left panels show representative scanning electron microscope (SEM) images of biofilms produced by the Xen31 strain 24 hours post-infection at 10 000x (top) and 20 000x (bottom) in wounds produced on the back of the experimental animal shown in the right-hand panel (see supplementary methods). Scale bars (lower left-hand corner) are 1 µm.

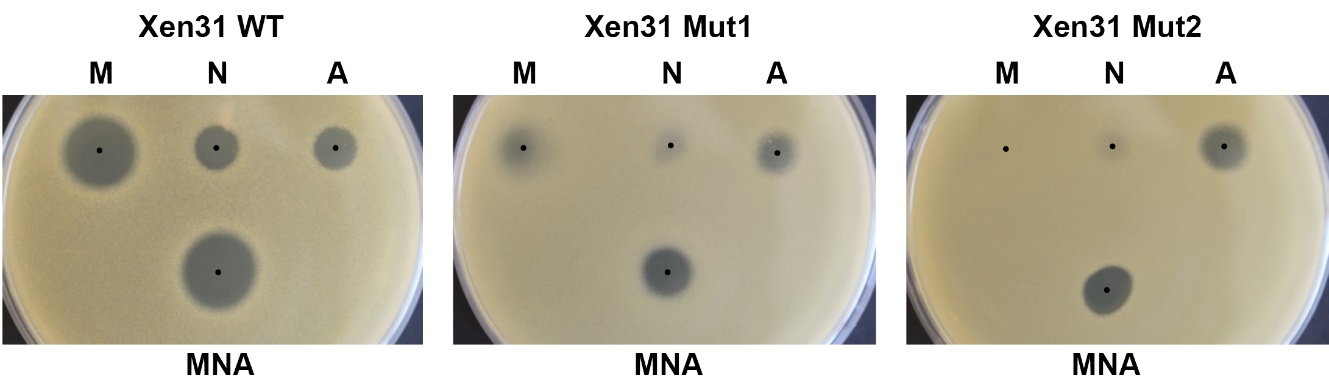

**Figure S9**. Susceptibility of Xen31 mutants recovered from wounded mice. Spot-on-lawn assay using MP1 (M), Nis (N), Aur (A) or the tricomponent combination (MNA) comparing Xen31 WT with two resistant isolates recovered at day 7 PI from the back of wounded mice.

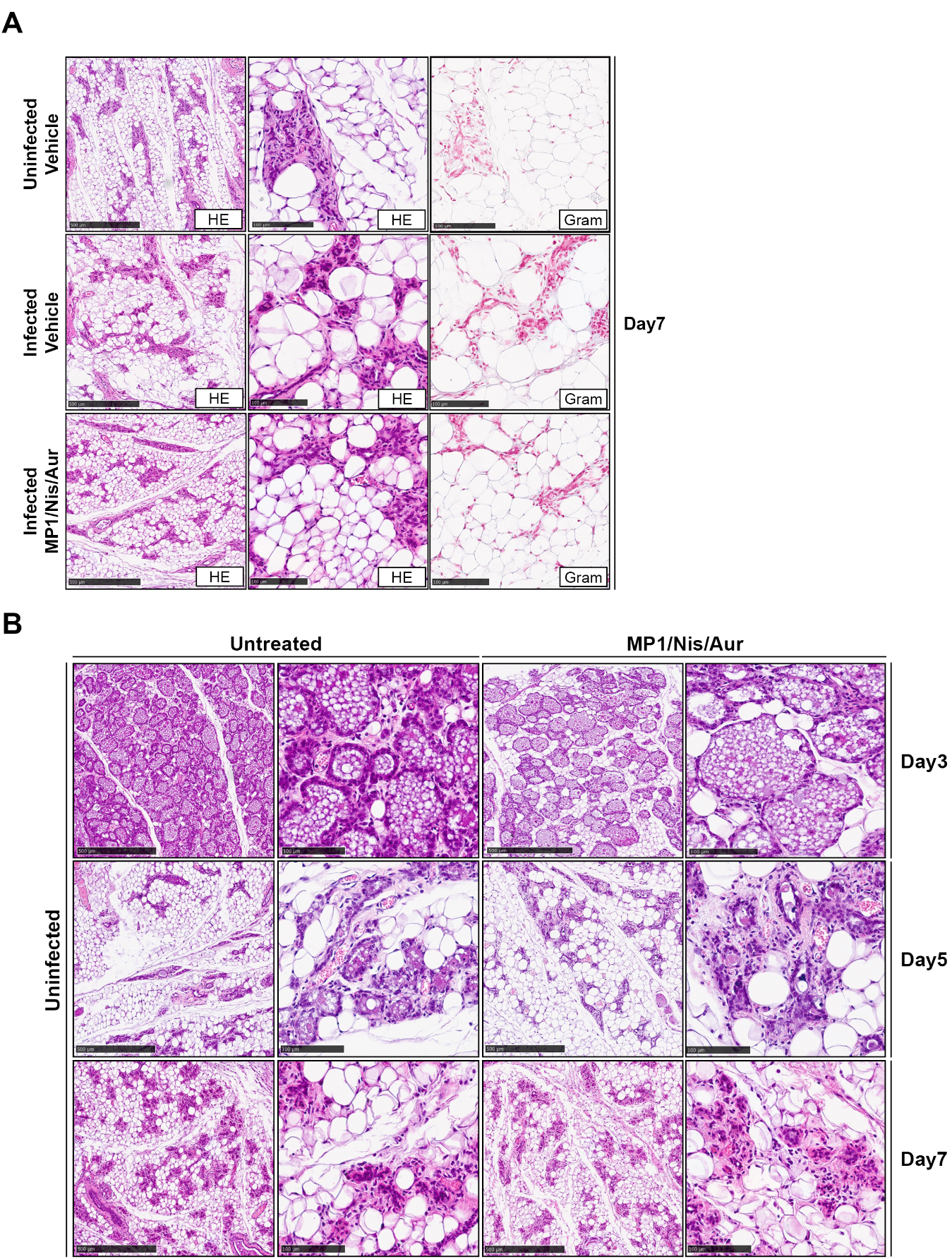

**Figure S10**. Pattern of infection at late time-points on the mastitis mouse model and effect of the micrococcin P1, nisin A and AuresinePlus combination on the mammary lobules*.* (a) Representative images of the histological examination of explanted mammary glands from uninfected mice injected with the antimicrobial vehicles, infected mice injected with the antimicrobial vehicles or infected mice injected with the antimicrobial combination at day 7 PI. Note that the antimicrobial treatment was stopped at day 5 PI. (b) Mammary glands of uninfected mice were injected with the MP1/Nis/Aur combination for the indicated time-points. All preparations in panel B were stained by H&E. In both panels, tissue section treatment and image representations are the same as those described in Figure 5.

**Supplementary Methods**

**Scanning electron microscopy of *S. aureus* in the wound**

Biopsies were fixed overnight in phosphate-buffered saline containing 3% glutaraldehyde. Fixed samplse was then dehydrated by successive washes in increasing ethanol concentrations (30, 50, 70, 90, 96%) for 10 min each, followed by four washes in absolute ethanol (4x10 min). Cells were dried by critical-point drying using a CPD 030 critical-point dryer (Bal-Tec) before sputter coating with palladium-gold using a Polaron Range sputter coater (Quorum Technologies). Microscopic examination was performed using an EVO50 EP scanning electron microscope (Zeiss).
